# Musleblind-1 regulates microtubule cytoskeleton in *C. elegans* mechanosensory neuron through tubulin mRNAs

**DOI:** 10.1101/2022.09.07.506915

**Authors:** Dharmendra Puri, Sarbani Samaddar, Sourav Banerjee, Anindya Ghosh-Roy

## Abstract

Regulation of microtubule cytoskeleton is fundamental for the development and maintenance of neuronal architecture. Recent studies have shown that regulated RNA processing is also critical for the establishment and maintenance of neural circuits. In a genetic screen using mechanosensory neurons of *C. elegans*, we identified a mutation in *muscleblind-1* as a suppressor of loss of kinesin-13 family microtubule destabilizing factor *klp-7*. Muscleblind-1(MBL-1) is an RNA-binding protein that regulates the splicing, localization, and stability of RNA. We found that *mbl-1* is required cell-autonomously for axon growth and synapse formation in the posterior lateral microtubule (PLM) neuron. Loss of *mbl-1* affects stability and plus-end-out organization of microtubules in the anterior process of PLM. These defects are also accompanied by abnormal axonal transport of the synaptic protein RAB-3 and loss of gentle touch sensation in *mbl-1* mutant. Our data showed that *mbl-1* is genetically epistatic to *mec-7* (β tubulin) and *mec-12* (a tubulin) for axon growth. The immunoprecipitation of MBL-1 pulls down the *mec-7, mec-12*, and *sad-1* mRNAs. Additionally, the *mbl-1* mutants show a reduction in the level and stability of *mec-7* and *mec-12* transcripts. Independently, *mbl-1* is epistatic to *sad-1* for synapse formation. Our work elucidated a previously unknown link between RNA binding protein and cytoskeletal machinery for the development and maintenance of the nervous system.

## Introduction

A highly ordered functional neuronal circuit comprises polarized nerve cells, which are compartmentalized into dendrites and axons that receive and transmit information unidirectionally. There are many reports which suggest that this structural and functional polarity of neurons is a function of cytoskeletal elements within the neuron (1-4). The cytoskeletal elements are regulated by intra- and extra-cellular signal transduction pathways during neuronal polarization (5-7). The organization of cytoskeletal component, microtubules, in the neuron, directs neuronal polarization, and development (8). In a vertebrate neuron, the axon has a plus-end out microtubule arrangement, facing towards the synapse, while in the case of dendrites, the microtubules are randomly oriented (8). In invertebrate dendrites, the microtubule arrangement is minus-end-out (9, 10). This polarized arrangement of microtubules is the basis for axonal transport, synaptic protein localization, and neurotransmitter release (8, 11).

Recent reports have identified the critical roles of RNA-binding proteins in neuronal development (12) and synaptic transmission (13-15). The disruption in these genes causes many neurological disorders (16-19). The Muscleblind-like protein family (MBNL) is an evolutionarily conserved RNA binding protein containing CCCH zinc-finger domains (20). MBNL regulates alternative splicing, alternative polyadenylation, mRNA localization, miRNA processing, and translation (21-23). The role of MBNL in the neural pathogenesis of myotonic dystrophy type 1(DM1), has been discussed in detail (24-27). In the mouse brain, loss of MBNL results in misregulated alternative splicing and polyadenylation causing defects in motivation, spatial learning, and abnormal REM(Rapid Eye Movement) sleep (21, 28-31). In mammals, MBNL family comprises MBNL1, MBNL2, and MBNL3 encoded by three different genes, and each gene has several isoforms (32). The functions of different isoforms of MBNL are different, which correlates with their differential localization (33-36). A recent report in *Drosophila* showed that Muscleblind (Mbl) is expressed in the nervous system and regulates alternative splicing of *Dscam2* for the development of the nervous system (37). Although there is an indication of a functional link between Muscleblind and microtubule cytoskeleton, (38), a comprehensive idea of how Muscleblind regulates microtubule cytoskeleton in neurons is unclear.

Mechanosensory neurons of *C. elegans*, responsible for gentle touch sensation, have been used to study microtubule regulation and neuronal polarization in vivo (39-41). In this study, we have identified a mutation in the *muscleblind-1* gene, as a suppressor of the touch neuron developmental defect in loss of kinesin-13 family microtubule depolymerase, *klp-7*. We have found that Musleblind-1(MBL-1) is required for axonal growth and synapse formation in the PLM touch neuron. Using live imaging of plus-end binding protein (EBP-2::GFP), and synaptic protein (GFP::RAB-3), we found that the microtubule stability is compromised in the absence of *mbl-1*, which leads to reduced vesicular transport. We further showed that *mbl-1* regulates mRNA stability of *mec-7* transcript and interacts epistatically with *mec-7* to control proper axon growth. Separately, *mbl-1* is epistatic to *sad-1* to control synapse formation in PLM neurons. Collectively, our data suggest that MBL-1 regulates cytoskeletal machinery for neuronal polarization by regulating mRNA stability.

## Results

### Loss of *muscleblind-1(mbl-1)* suppresses the multiple axon-like projections phenotype due to loss of *klp-7* in touch neurons

In *C. elegans*, six mechanosensory neurons are responsible for gentle touch sensation. The anterior neurons are known as Anterior Lateral Microtubule (ALM) and posteriors are known as Posterior Lateral Microtubule (PLM) (white arrowheads, Figure 1A). ALM and PLM neurons grow their axons laterally towards the anterior side and make a connection to their respective postsynaptic neurons through a ventral synaptic branch (white arrows, Figure 1A). Additionally, PLM also has a short posterior process (double-sided white arrow, Figure 1A). Recently, we showed that loss of kinesin-13/KLP-7 microtubule depolymerizing protein, leads to multiple axon-like phenotype in ALM neuron (orange arrow, Figure 1A), and the overgrowth phenotype of the PLM posterior process (double-sided white arrow, Figure1A),Figure 1A) due to excessive stabilization of the microtubule cytoskeleton (7). Destabilization of microtubules using colchicine or loss of tubulin subunits suppresses the axon overgrowth phenotype seen in the *klp-7* mutant (7). Therefore, we hypothesized that a suppressor screen for the neuronal phenotype in the *klp-7* mutant background, might help identify pathways that regulate microtubule cytoskeleton in the neuron. One of the suppressors, *ju1128*, that suppressed the ectopic extension phenotype of ALM neuron (red arrow, Figure 1A, C), maps to the locus of the *mbl-1* gene. Several lines of evidence support that *ju1128* is an allele of *mbl-1* gene that codes for the RNA binding protein Muscleblind-1/MBL-1. First, the recombination cross with the Hawaiian strain followed by restriction fragment length polymorphism (RFLP) analysis (42, 43) of the F2 progenies indicated an association of the suppression with the right arm of the X chromosome (red arrowhead, Figure S1A). The whole-genome sequencing of the outcrossed and re-isolated suppressor (44) revealed a strong peak (frequency of pure parental alleles) at the precise location of the *mbl-1* gene (Figure S1B). After annotating the SNPs obtained from the Cloudmap analysis of whole-genome sequencing data (44), we identified a C-T transition at the 17002646^th^ base pair position of chromosome X (Figure 1B). The 87^th^ nucleotide of the third exon of the *mbl-1* gene is mutated in *ju1128*, which introduces a premature stop codon in place of glutamine amino acid.

**Figure 1:**
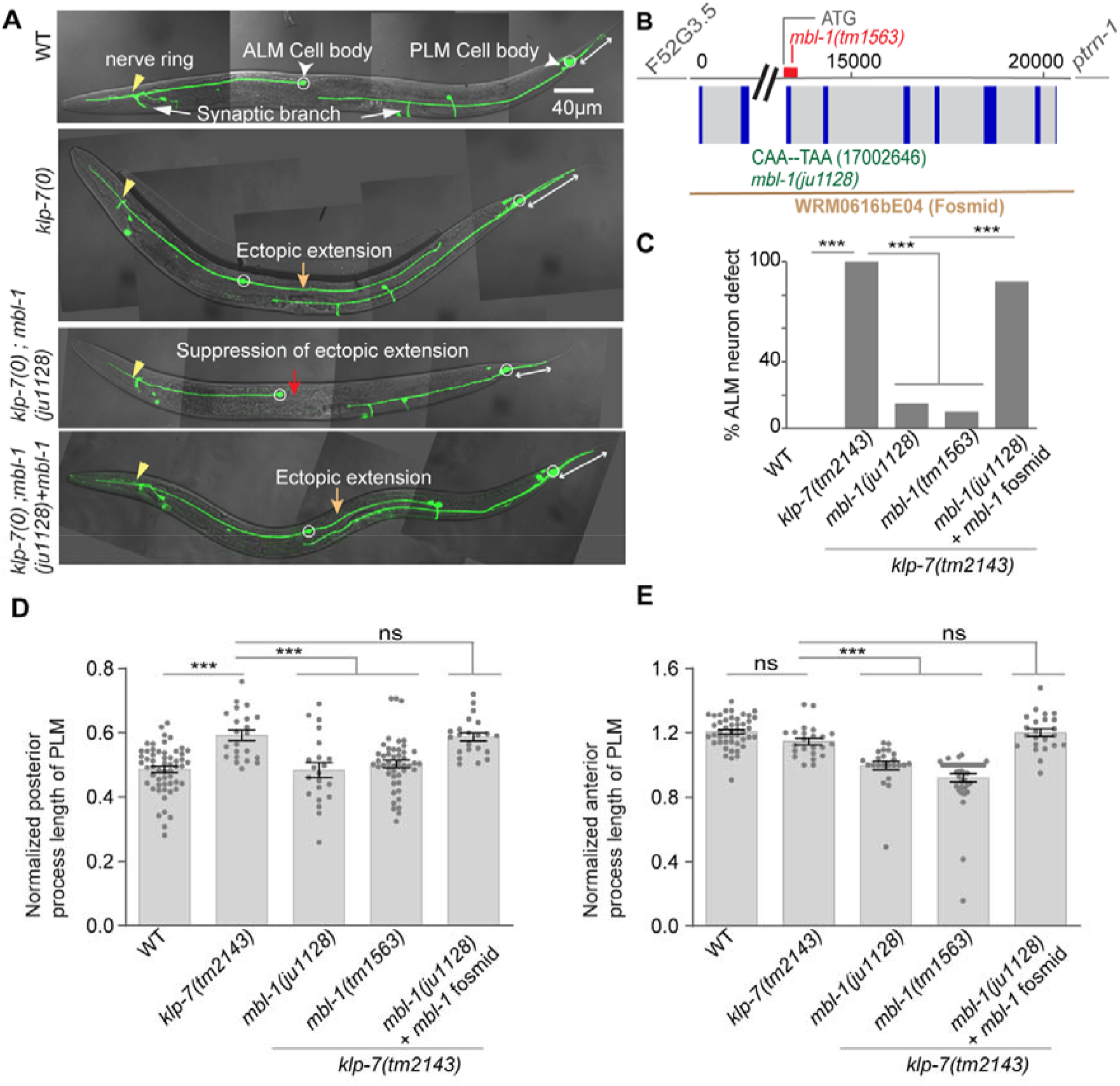
Characterization and mapping of *ju1128* mutation in *mbl-1*. (A) Confocal images of ALM and PLM neurons of a wild-type, *klp-7(0)* and suppressor *klp-7(0); ju1128* expressing *muIs32* (*Pmec-7*::GFP) at larval-stage four (L4) animal. The ectopic extension of the ALM posterior process in *klp-7(0)* is marked by a yellow arrow while PLM posterior process is marked by a double sided white arrow. The suppression of *klp-7(0)* ectopic extension in suppressor *klp-7(0); ju1128* marked by red arrow. (B) The schematic of the exon and intron of *mbl-1* gene and the genetic position of *ju1128* and *tm1563* alleles with respect to the *mbl-1* locus. The fosmid WRM0616bE04, which exclusively covers the *mbl-1* gene is also shown. (C) Quantification of suppression of ectopic extension of ALM in *klp-7(0); ju1128* background and the rescue of *klp-7(0)* ectopic extension of ALM in *klp-7(0); ju1128+* WRM0616bE04 fosmid background. N = 3-5 independent replicates, n (number of neurons) = 100-150. (D-E) The normalized length of posterior (D) and anterior process (E) of PLM, in *klp-7(0)*, suppressor *klp-7(0) ju1128*, and *klp-7(0); ju1128+ WRM0616bE04* fosmid background. Normalized length = (Actual length/distance between the PLM cell body and vulva for the anterior process and the distance between the PLM cell body to the tip of the tail for the posterior process). N = 3-4 independent replicates, n (number of neurons) = 21-47. For C, ***P<0.001; Fisher’s exact test. For D-E, ***P<0.001; ANOVA with Tukey’s multiple comparison test. Error bars represent SEM, ns, not significant.

The mapping results were further confirmed by expressing the fosmid WRM0616bE04, in the background of *klp-7(0); ju1228*. This fosmid exclusively contains the complete locus of *mbl-1* gene. The extrachromosomal expression of *mbl-1* strongly rescued the suppression of the multipolar phenotype in the ALM neuron (yellow arrow, Figure 1A, C) and also rescued the overgrowth phenotype of the posterior process in the PLM neuron (double-sided white arrow, Figure 1A, D-E). Another allele of *mbl-1, tm1563*, which is a deletion mutation in the third exon (Figure 1B), also suppressed the multiple axon-like phenotypes in *klp-7(0)* (Figure 1C). These observations suggested that *ju1128* is a mutation in the *mbl-1* gene, and the loss of function of *mbl-1* suppresses the neuronal phenotypes in *klp-7(0)*.

### Muscleblind-1(MBL-1) regulates axon growth and synapse formation in PLM neuron

To understand the role of *mbl-1* gene in touch neuron development, we removed the *klp-7(tm2143)* allele from the suppressor background. In the wild-type, PLM axon crosses the vulva and approaches the ALM cell body anteriorly (magenta arrowhead, Figure 2A), making a ventral synaptic branch the PLM axon makes a synapse near the vulva (white arrow, Figure 2A). However, in both the mutant alleles of *mbl-1*, PLM axon terminates before the vulva or at the vulval position (yellow arrow, Figure 2A), which we termed as ‘short neurite’ phenotype. Nearly 88% of the PLM neurites are short in *mbl-1* mutants (Figure 2A-B). Since the Muscleblind-1 protein is known to play its role both in muscles (20) and in neurons (37, 45), we wanted to know the tissue-specificity of function of *mbl-1* in neurite development. When we expressed *mbl-1* cDNA in muscles using *Pmyo-3::mbl-1*cDNA in the *mbl-1(0)*, we did not see any rescue of the ‘short neurite’ phenotype (Figure 2B). However, when we expressed *mbl-1* cDNA either pan-neuronally using *Prgef-1::mbl-1* or only in the mechanosensory neurons using Pmec*-4::mbl-1* in *mbl-1(0)* background, we saw a significant rescue of the short neurite phenotype, which was comparable to the rescue obtained using the *mbl-1* genomic fragment (Figure 2A-B).

**Figure 2:**
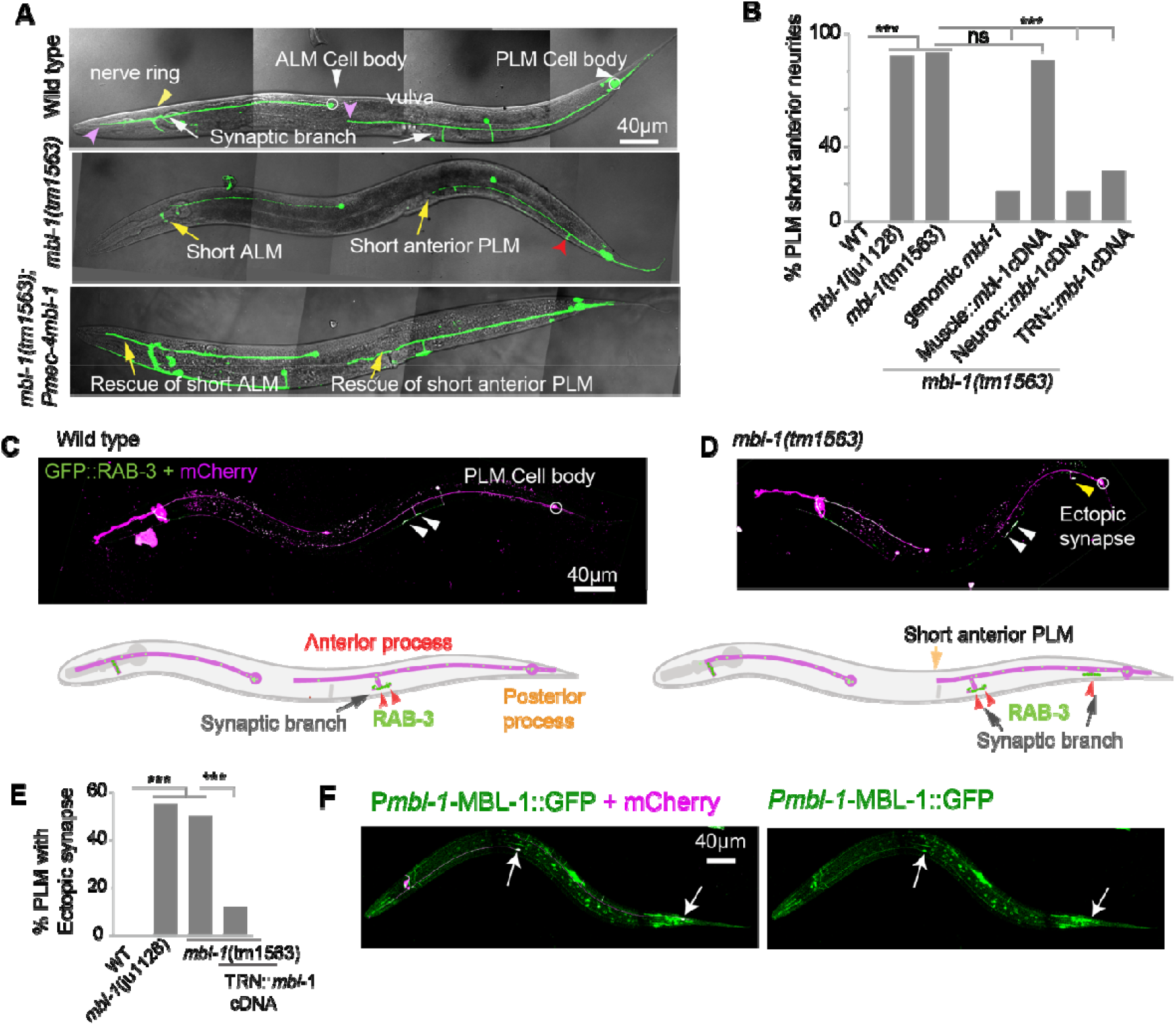
*mbl-1* mutant display defect in axon growth, and synapse formation in PLM neuron. (A) Confocal images of ALM and PLM neurons at larval four-stage (L4) in WT, *mbl-1(tm1563)*, and *mbl-1(tm1563)*; *Pmec-4::mbl-1* background. The ends of ALM and PLM anterior neurites are marked with a magenta arrowhead. *In mbl-1(tm1563)* short anterior process of ALM and PLM is marked by yellow arrow and the rescue of the short anterior process in *mbl-1(tm1563); Pmec-4::mbl-1* is also marked by yellow arrow. The presence of ectopic synapse in *mbl-1(tm1563)* is marked by a red arrowhead. (B) Quantification of short PLM anterior neurite at L4 stage in two alleles of *mbl-1(0)* and different rescue backgrounds. N = 4-5 independent replicates, n (number of neurons) = 100-200. (C and D) Image and schematic of synapse in PLM in both wild-type (C) and *mbl-1(tm1563)* (D) at the L4 stage. PLM Synapse (white arrowheads) was visualized with the *Pmec-7*::GFP::RAB-3 transgene. (D) The presence of an ectopic synapse in the *mbl-1(tm1563)* background is marked by a yellow arrowhead. (E) Quantification of ectopic synapse phenotype in *mbl-1(tm1563)* and *mbl-1(tm1563); Pmec-4::mbl-1* backgrounds. N = 3-4 independent replicates, n (number of neurons) = 80-100. (F) Confocal images of ALM and PLM neurons (arrows) in worms expressing *Pmbl-1::*MBL-1::GFP (*wgIs664*) in P*mec-4:mCherry* (*tbIs222*) background.

It is known that the PLM axon makes physical contact with the BDU interneuron through a gap junction (46). In the *mbl-1(0)* mutant, the physical contact between the PLM axon and the BDU interneuron is lost as the PLM axon is short (Figure S2A-C). Using the GAP junction reporter UNC-9::GFP (46) in the *mbl-1(0)*, we found that the GAP junction at the tip of the PLM anterior neuron was missing (Figure S2D-F). The ALM axon, in the wild-type, makes a synapse at the nerve ring (Figure 2A) and extends anteriorly terminating near the tip of the head (magenta arrowhead, Figure 2A). In *mbl-1(0)*, the ALM terminates at the nerve ring itself (yellow arrow, Figure 2A).

In addition to the short neurite phenotype, *mbl-1(0)* also had an ectopic synapse near the PLM cell body (red arrowhead, Figure 2A). We used a presynaptic marker (*Pmec-7*::GFP::RAB-3) (47) to characterize this ectopic synapse. In the wild-type, PLM anterior process makes a synapse near the vulva (Figure 2C white arrowhead), while in the *mbl-1(0)* we observed an ectopic synapse near the cell body (yellow arrowhead, Figure 2D) in addition to the original synapse (white arrowhead, Figure 2D). In both mutant alleles of *mbl-1*, we observed nearly 50% of the PLMs showing ectopic synapses (Figure 2E), and this phenotype was rescued by extrachromosomal expression of *mbl-1* cDNA in mechanosensory neurons (Figure 2E). We used another presynaptic active zone marker ELKS-1 (ELKS-1 :: TagRFP) (48), which also showed ectopic synapse near the PLM cell body, similar to GFP: RAB-3 (Figure S2G-I). Next, we checked the localization of MBL-1 using *pmbl-1*::MBL::GFP. MBL-1::GFP was highly enriched in many neurons including ALM and PLM touch neurons (Figure 2F).

To determine whether the mutation in the *mbl-1* gene affects other classes of neurons, we visualized the D-type GABAergic motor neurons using *Punc-25*::GFP reporter transgene. In the *mbl-1(0)* mutant, we noticed that on the dorsal side, there are often gaps (red arrowhead, Figure S2J-L) indicating a synaptic defect, as seen in the case of DA9 neuron in *mbl-1* mutant previously (45). All these observations suggest that the *mbl-1* gene is required for neurite growth and synapse formation in neurons.

However, we did not observe any noticeable morphological defect in the loss of function mutants of MEC-8/RBMPS, MSI-1/MSI-2, UNC-75/CELF5, and EXC-7/ELAVL4, the other four known RNA binding proteins, in ALM and PLM neurons (Figure S2 M-O).

### MBL-1 regulates microtubule polarity and stability in the anterior process of PLM neuron

Since the *mbl-1* mutant suppressed the neuronal overgrowth phenotype caused due to stabilization of microtubules in the *klp-7* mutant, we looked for any possible defects in the microtubule cytoskeleton in *mbl-1(0)*. We did time-lapse live imaging of EBP-2:GFP, (9, 49). We determined the polarity of the microtubules from the direction of microtubule growth from the plus ends, as seen in the kymographs (Figure 3B) from the regions of interest in the anterior and posterior processes of PLM neuron (Figure 3A). In the wild-type background, the PLM anterior process had the majority of the EBP-2::GFP movements away from the cell body (plus-end-out, green trace), while in the posterior process, EBP-2::GFP movements were seen both away from and towards the cell body (minus-end-out, magenta trace, Figure 3A-B), as reported before (7). We plotted the fraction of microtubule tracks with ‘plus-end-out’ or ‘minus-end-out’ orientation (Figure 3C). In the *mbl-1(0)*, the % of the microtubules with plus-end-out orientation is significantly decreased in the anterior process (Figure 3C). In the posterior process, the microtubule arrangement was similar to wild-type (Figure 3C). In addition, we noticed that in the *mbl-1(0)*, the number of EBP-2::GFP tracks was higher in the anterior and posterior process of PLM (Figure 3D) as compared to the wild-type. The growth length and duration of these tracks were significantly smaller in the PLM anterior process as compared to wild type, while in the posterior process the growth duration is smaller than wild type. (Figure 3 E-F). These observations suggest that *mbl-1(0)* has increased microtubule dynamics. When we compared the frequency distribution of microtubule polarity in PLM processes, we found that in *mbl-1(0)* PLM anterior had a nearly normal distribution with a mode value of 0.5, while in wild-type mode value was 1.0 (Figure S3A-B) However, in the PLM posterior process of *mbl-1(0)* mode value was nearly 0.5, similar to the wild-type (Figure S3 C-D). We classified the PLM processes with a fraction polarity value of 0.8 or more as a unipolar process (gray shaded box, Figure S3A-D). A PLM process with a fraction polarity value lower than 0.8 was categorized as a process with a mixed microtubule arrangement. Based on this criterion, in the *mbl-1(0)* background, 80 % of the PLM anterior processes were with mixed microtubule polarity, while in the wild type, 13 % of the anterior processes were mixed (Figure 3G). All these observations suggest that the loss of *mbl-1(0)* affects both microtubule dynamics as well as arrangement in PLM neuron.

**Figure 3:**
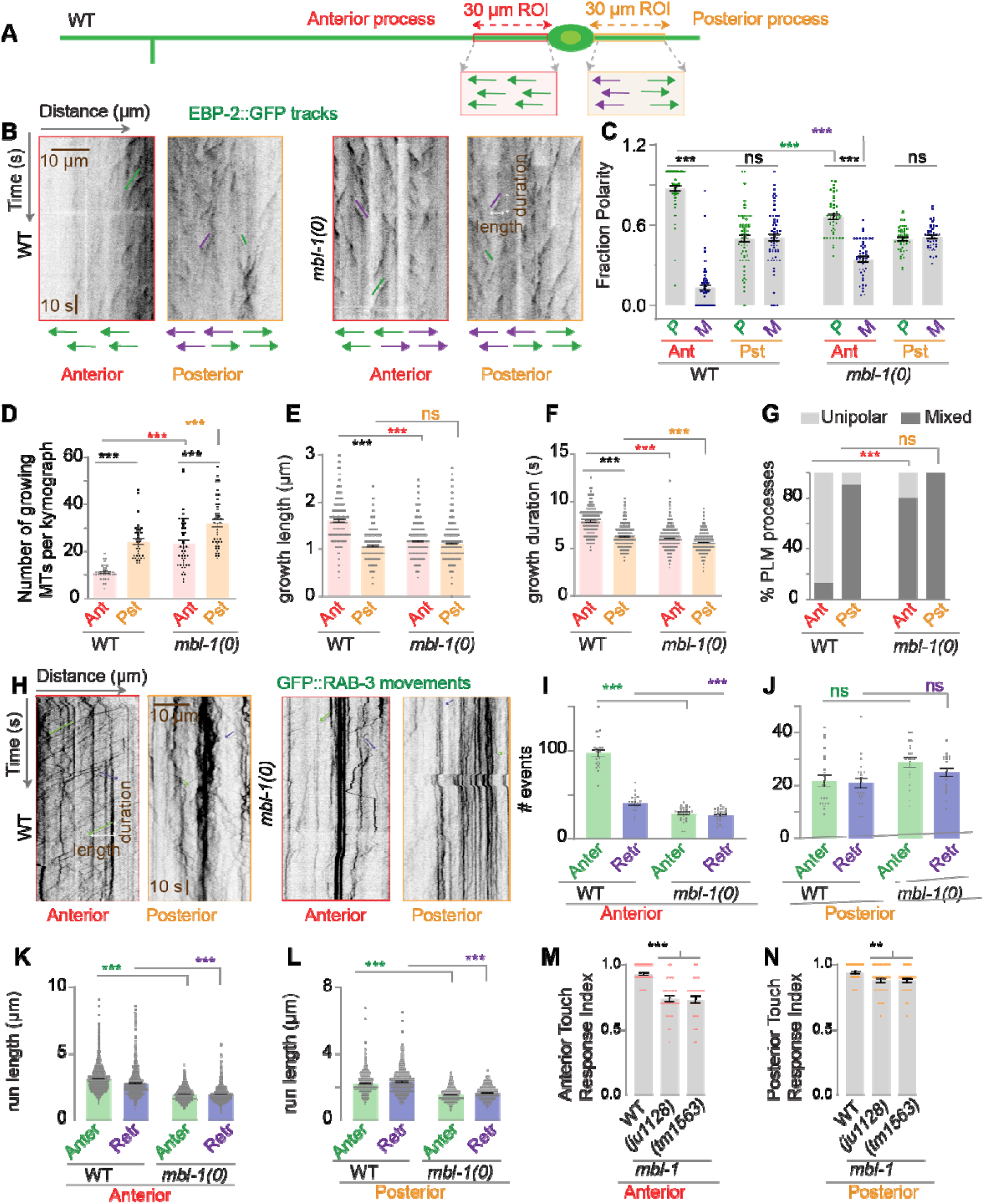
*mbl-1* mutant affects microtubule dynamics in PLM neuron. (A) Schematic of the PLM neuron. 30µm Regions of interest (ROIs), marked in red and orange for anterior and posterior processes respectively, used for the analysis of time-lapse movies of *Pmec-4*::EBP-2::GFP (*juIs338*). (B) Representative kymographs of EBP-2::GFP obtained from the above-mentioned ROIs in anterior and posterior processes of PLM in wild type and *mbl-1(0)* background. The green and magenta traces on kymographs represent microtubule growth events away from the cell body (Plus end out) and towards the cell body (Minus end out), respectively. (C) The histogram is showing the fraction of microtubules with plus-end-out’ (P) or ‘minus-end-out’ (M) polarity in wild-type and *mbl-1(0)* in PLM anterior and posterior processes. N=3-5 independent replicates, n (number of worms) = 44-62. (D) The histogram represents the number of growing microtubules in PLM anterior and posterior processes in wild-type and *mbl-1(0)*. N=3-5 independent replicates, n (number of worms) = 36-46. (E and F) Growth length (E) and growth duration (F) of the tracks, measured from net pixel shift in the X and Y axis respectively, from kymographs shown in B. N=3-5 independent replicates, n (number of tracks) = 227-1074. (G) % of PLM processes with microtubules organized either in unipolar or mixed arrangement manner, N=3-5 independent replicates, n (number of worms) = 36-46. (H) Representative kymographs of time-lapse movies of *Pmec-7*::GFP::RAB-3 (*juIs821*) as obtained from the above-mentioned ROIs (A) of anterior and posterior processes of PLM, in wild type, and *mbl-1(0)* background. The green and magenta traces on kymographs represent anterograde (green trace) and retrograde (magenta trace) movement events away from the cell body (anterograde) and towards the cell body (retrograde), respectively. (I and J) Quantification of the number of anterograde (Anter) and retrograde (Retr) movement events of GFP::RAB-3 particles obtained from kymographs (H) in PLM anterior (I) and posterior (J) processes in wild-type and *mbl-1(0)*, N=3-5 independent replicates, n (number of worms) = 21-32. (K and L) run length in PLM anterior (K) and posterior (L) processes in wild-type and *mbl-1(0)*, measured from net pixel shift in X-axis direction as shown in the kymograph (H). N=3-5 independent replicates, n (number of tracks) = 456-2131. (M and N) The histogram shows the anterior (M) and posterior (N) gentle touch response index of the worm in the wild-type and two alleles of the *mbl-1* gene. N=3 independent replicates, n (number of worms) =31-50. For C -F, and I-N ***P <0.001; ANOVA with Tukey’s multiple comparison test. For G, ***P <0.001. Fisher’s exact test. Error bars represent SEM. ns, not significant.

Since neuronal microtubule organization and stability is affected due to loss of *mbl-1*, we checked vesicular transport in *mbl-1(0)* using a GFP reporter of presynaptic protein RAB-3 (47). We imaged GFP::RAB-3 in similar ROIs (Figure 3A), which were used for imaging EBP-2::GFP. In the *mbl-1(0)*, the number of anterograde and retrograde transport events were reduced in the PLM anterior process as compared to the wild type (Figure 3H, I). However, in the PLM posterior process, the transport events were similar to the wild type (Figure 3 H, J). We also noticed that in the anterior process, most of the particles were static (Figure 3H) in the *mbl-1(0)*. The run length of the transport event was less in the *mbl-1(0)* as compared to the wild type (Figure 3 K, L).

It has been seen that the abnormal arrangement of microtubules in mechanosensory neurons leads to defects in gentle touch sensation (39). In the *mbl-1(0)* background, the anterior and posterior touch response was significantly reduced in both the alleles of the *mbl-1* (Figure 3 M-N).

Collectively, our findings suggest that, within PLM axons, loss of MBL-1 results in microtubules having mixed polarity and enhanced dynamicity; which consequently leads to defects in axonal transport.

### *mbl-1* genetically interacts with the cytoskeletal components and its regulatory genes to control neurite growth of PLM neuron

MBL-1 is a Zinc finger family RNA binding protein that preferentially binds to the CGCU sequence of target RNA (50) (Figure S4A). Using the oRNAment database (http://rnabiology.ircm.qc.ca/oRNAment) (51), from a pool of mRNAs expressed in the PLM neuron as reported in the CeNGEN database (52), we filtered the ones with a binding site for MBL-1. We found 2000 such targets of MBL-1 in PLM neurons (Figure 4A) (Table S1). From this pool of genes, using gene ontology (GO) analysis, we short-listed four sets of genes for further analysis, based on their involvement in (1) Microtubule-based processes, (2) Axon development, (3) Regulation of synapse structure, and (4) Axodendritic transport (Figure 4A).

**Figure 4:**
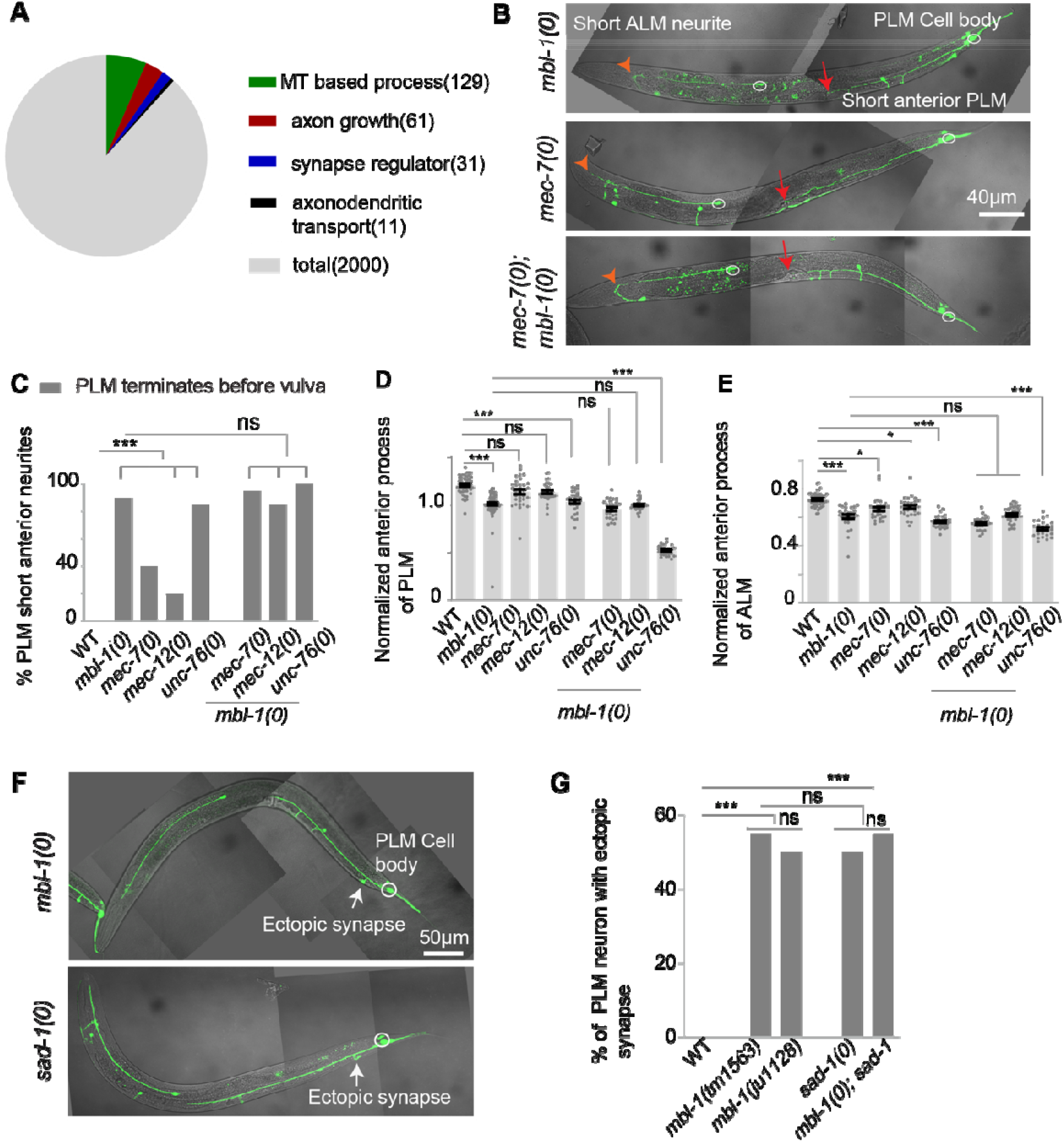
*mbl-1* genetically interacts with mutants affecting microtubule cytoskeleton and *sad-1* kinase for proper axon growth and synapse formation in the PLM neuron. (A) Gene ontology (GO) analysis of putative targets of MBL-1 (transcripts with an MBL-1 binding site) expressed in PLM neuron (B) Representative confocal images of *mbl-1(0), mec-7(0)* and *mbl-1(0) mec-7(0)*. The red arrow indicates a short anterior process in PLM while the orange arrowhead indicates a short ALM anterior process. (C) The histogram is showing the percentage of PLM neurons with a short anterior process. (D) The normalized length of anterior process of PLM neuron in different backgrounds. (E) Quantification of the normalized length of anterior process of ALM neurons. For ALM neuron, normalized length = (Actual length of ALM anterior process /distance between vulva and the tip of the nose). (F-G) The representative confocal images (F) and quantification (G) of ectopic synapses (marked with gray arrow) in the *mbl-1(0)* and *sad-1(0)* backgrounds. For C-F, and G independent replicates (N) = 3-5 and the number of neurons (n) = 30-150. For C and G; ***P<0.001. Fisher’s exact test. For D-E ***P <0.001; ANOVA with Tukey’s multiple comparison test. Error bars represent SEM, ns, not significant

In the microtubule-based processes, we got 129 genes (Table S2) (Figure 4A) out of which we tested 13 genes (Table S3), that are either a part of microtubule structure or are regulators of microtubule dynamics, for example, touch neuron-specific tubulins (*mec-7* and *mec-12*) and CRMP-2 (*unc-33*) which helps in microtubule polymerization (41, 53). In the axon growth-related genes category, we got 61 genes (Table S2) (Figure 4A), and we tested 16 of these genes (Table S3), which are known to play a role in axon growth, for example, *unc-51* and *unc-53* (54-56).

In the regulators of synapse structure category, we got 31 genes (Table S2) (Figure 4A), out of which we checked 5 genes (Table S3) that are known to regulate synapse development, for example, SAD-1 kinase (5, 6).

In the axodendritic transport-related genes category, we found 11 genes (Table S2) (Figure 4A) and we have tested 4 genes (Table S3) that are known to regulate motor-based transport such as *unc-104* (57, 58).

Additionally, a recent report identified a set of 235 genes that are downregulated in the *mbl-1* mutant (59). This set of genes was again sorted using GO analysis to short-list candidates from above mentioned categories. It gave us *mec-7, mec-12*, and *klp-13* [, that are involved in microtubule-based processes. However, from this set of genes, we could not find any candidates linked to axon development, synapse structure, or axodendritic transport.

Next, we phenotyped the mutants of the above-mentioned candidate genes, for a short neurite or ectopic synapse in the PLM neuron, as observed in the *mbl-1(0)* mutant. We found that mutations in the tubulin genes *mec-7 (*β tubulin*), mec-12 (*α tubulin), and vesicular adaptor protein (*unc-76)* lead to a short neurite phenotype in the PLM anterior process (red arrow, Figure 4B-E), similar to *mbl-1(0)*. The transcript of each of these genes has an MBL-1 binding site (Figure S4 A-C).

In the *mbl-1(0) mec-7(0)* double mutant the length of the PLM axon remained the same as that observed in *mbl-1(0)* single mutant, showing that the phenotype is not additive (Figure 4B-E). However, in *mbl-1(0); unc-76(0)* double mutant, the length of the PLM axon was shorter than either of the single mutants (Figure 4D). These results suggest that *mbl-1* and *mec-7* are genetically working in the same pathway, while *mbl-1* and *unc-76* might be genetically working in a parallel pathway, as they showed a synergistic effect. A similar observation was also made in ALM neurons (light red arrowhead, Figure 4B; E).

The transcript of the *sad-1* kinase is a known target of MBL-1 in touch neurons (60) (S4D). We found that the loss of function mutant of *sad-1* has a synapse defect similar to that of *mbl-1* mutant (grey arrow, Figure 4F-G). And in the *sad-1(0); mbl-1(0)* double mutant the extent of ectopic synapse defect is the same as in single mutants (Figure 4G). These results suggest that *mbl-1* and *sad-1* are genetically working in the same pathway for the formation of the synapse at the correct location in PLM neuron. This observation is an indication that MBL-1 might be regulating *sad-1* to regulate proper synapse targeting in touch neurons.

### MBL-1 regulates the stability of *mec-7* and *mec-12* mRNAs

MBL-1 protein is well documented to function in alternative splicing, stability, and localization of RNAs (21-23). To understand how the transcripts of the short listed target genes are affected in *mbl-1* mutant, we performed quantitative RT-PCR analysis *mec-7, mec-12, unc-76*, and *sad-1* in wild-type and *mbl-1(0)*. We did not observe any changes in the lengths of the PCR products of these genes in *mbl-1(0)*, which was verified using multiple primers (Figure 5A-B, Figure S5A-B). However, there was a reduction in total transcript of *mec-7* and *mec-12* in the *mbl-1(0)* mutant (Figure 5A-B). Such a reduction in the total transcript was not noticed in the case of control *aak-2* or *tba-1* genes (Figure 5A-B). We found a 0.4717± 0.03720 fold (mean ± SEM) (2−ΔΔCT) decrease in total *mec-7* levels (***P = 0.0005; ANOVA, Tukey’s Multiple Comparison Test) in the *mbl-1(0)* background as compared to control (Figure 5B-C, S5C). Similarly, there was a significant decrease in *mec-12* levels in *mbl-1(0)* (Figure 5B-C, S5 C). However, we did not observe any changes in the transcript length or the amount of *unc-76* and *sad-1* in the *mbl-1(0)* background (Figure S5B-C, Figure 5C). This led to the conclusion that MBL-1 regulates the total transcript levels of *mec-7* and *mec-12* genes.

**Figure 5:**
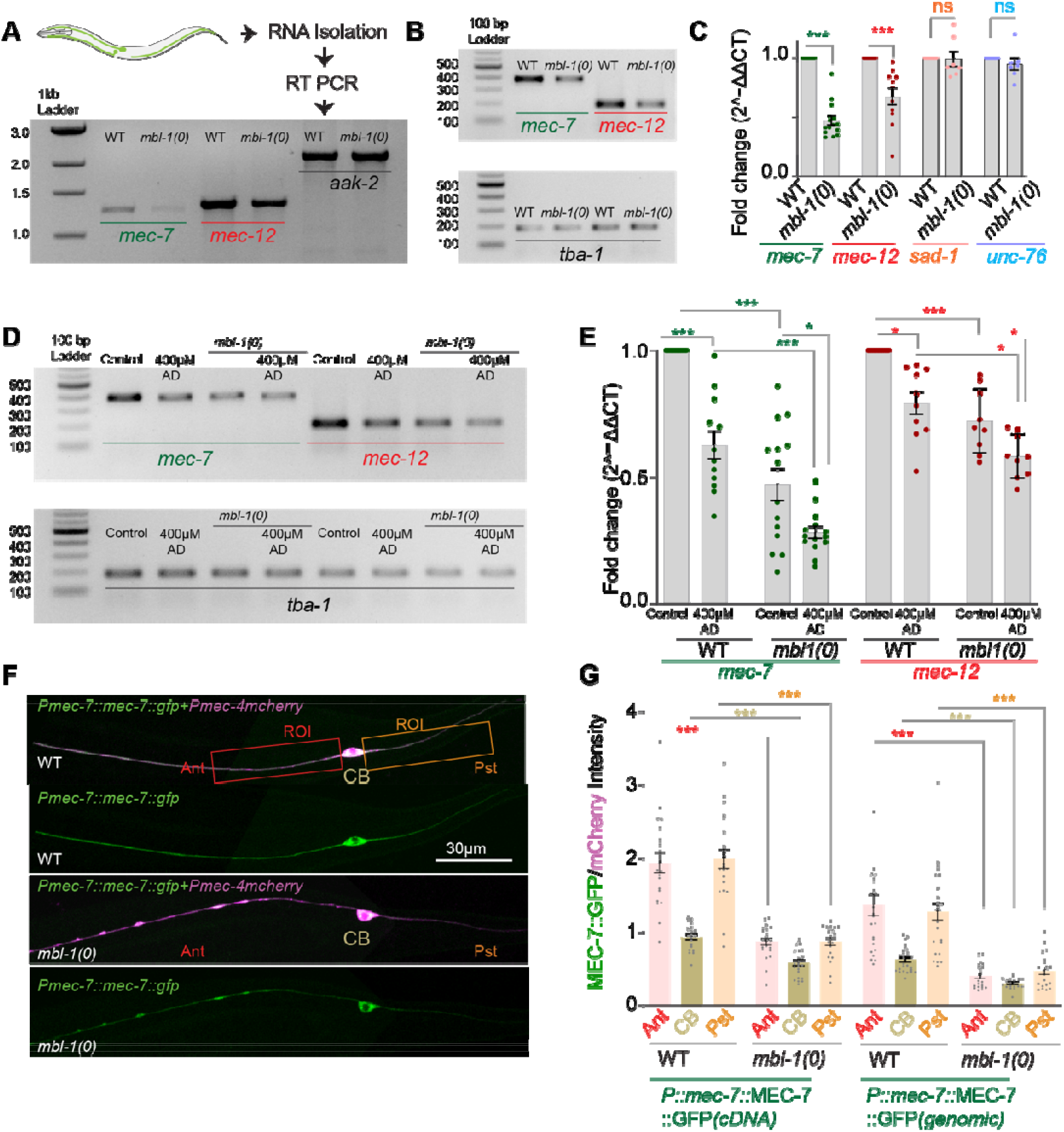
MBL-1 regulates the level of *mec-7* and *mec-12* tubulin transcripts. (A) Illustration depicting the method for reverse transcription (RT)-PCR and representative agarose gel image showing transcript length of *mec-7, mec-12*, and *aak-2* (control) in the wild-type and *mbl-1(0)* background. (B) Representative agarose gel image from the sample of quantitative real-time (qRT) PCR, showing a reduction in transcript amount in *mec-7* and *mec-12* in the wild-type and *mbl-1(0)* background. *tba-1* has been used as an internal control. (C) Quantification of qRT-PCR of the transcripts of *mec-7, mec-12, sad-1*, and *unc-76* in the wild-type and *mbl-1(0)* background. Independent replicates (N) = 10-11 and the number of reactions (n) = 11-15. (D-E) Representative agarose gel (D) and quantification of transcript (E*) mec-7* and *mec-12* in the 400μM actinomycin D treated worms in the wild-type and *mbl-1(0)* backgrounds. Independent replicates (N) = 9-10 and the number of reactions (n) = 9-14. (F-G) Representative confocal images of the worms (F) expressing *Pmec-7::mec-7::gfp (shrEx473)* and *Pmec-4::mCherry (tbIs222)* and (G) quantification of ratio (*gfp/ mCherry)* of intensity from 50 μm regions of interest(ROI) in the anterior (Ant) and posterior (Pst) process of PLM and also from the PLM cell body (CB). independent replicates (N) = 3-4 and the number of neurons (n) = 20-25. For C, E, and G *, P < 0.05; ***, P < 0.001; ANOVA with Tukey’s multiple comparison test. Error bars represent SEM.

We hypothesized that the total mRNA transcript is a quantitative measure of MBL-1 regulating either the transcription of these genes or the stability of the mRNAs. To test whether MBL-1 regulates transcription or stability of its target mRNAs, we fed the worms Actinomycin D (Act D), a transcription inhibitor (61). Act D treatment would block the expression of new transcripts, providing a suitable context to study whether any change occurs to the pre-existing *mec-7* and *mec-12* transcripts in *mbl-1(0)* mutants. As expected, the wild-type worms fed on 400 µM Actinomycin D, showed a 0.6264 ± 0.05498-fold reduction (mean ± SEM) (2−ΔΔCT) in the levels of *mec-7* transcripts (Figure 5D-E). A similar trend was also observed in the *mec-12* transcript levels of the wild type (Figure 5D-E). These results indicated that Act D could successfully block transcription in wild-type worms. When the *mbl-1(0)* mutant was grown on 400 µM Actinomycin D, we observed a further reduction in the levels of *mec-7* and *mec-12* transcript as compared to the wild-type (Figure 5D-E), grown on 400 µM Actinomycin D. This reduction observed in the *mbl-1(0)* mutants as compared to the wild type is interesting, as it illustrates the fate of the pre-existing transcripts in the absence of MBL-1 and in the absence of any new transcription. As transcription was blocked, the observed additional decrease could be attributed to the increased instability of the pre-existing transcripts in the absence of MBL-1.

We further validated this reduction in the amount of *mec-7* transcript using a translational reporter. We observed a diminished absolute as well as normalized (MEC-7::GFP/ mCherry) MEC-7::GFP intensity in the anterior and the posterior process of PLM neurons, as well as in the cell body (Figure 5F-G, S6A-B). From these results, we concluded that MBL-1 is regulating the stability of *mec-7* and *mec-12* mRNA in PLM neurons.

### MBL-1 interacts with *mec-7* and *sad-1* mRNA in the mechanosensory neuron

To ascertain that *mec-7*, and *mec-12* mRNAs interact with MBL-1 protein, we performed Ribonucleoprotein Immuno-Precipitation (RIP). We pulled down RNA-bound MBL-1, from transgenic wild-type worms, ubiquitously expressing MBL-1::GFP::FLAG under its native promoter, using an anti-Flag antibody. (Figure 6A). We detected an enrichment of *mec-7, mec-12*, and *sad-1* transcripts in the immuno-precipitated (IP) sample as compared to the control sample (Figure 6 B-C). However, we did not find *unc-76* mRNA enrichment upon immunoprecipitation of MBL-1 from these worms (Figure 6 B-C). We used worms expressing KLP-7 :: GFP::FLAG under its native promoter (62) as a control sample for this experiment. To substantiate further that MBL-1 protein is associated with *mec-7* mRNAs in the mechanosensory neurons, we immuno-precipitated GFP from *mbl-1(0)* worms expressing MBL-1::GFP in the mechanosensory neurons under the *Pmec-4* promoter. The *Pmec-*4::MBL-1::GFP transgene could rescue nearly 75% of *mbl-1* loss of function phenotype. We found an enrichment of *mec-7* and *sad-1* mRNA in the Immuno-Precipitation (IP) sample in contrast to the control sample (Figure 6 D-E). From these results, we concluded that MBL-1 specifically associates with *mec-7* and *sad-1* mRNA in the mechanosensory neurons.

**Figure 6:**
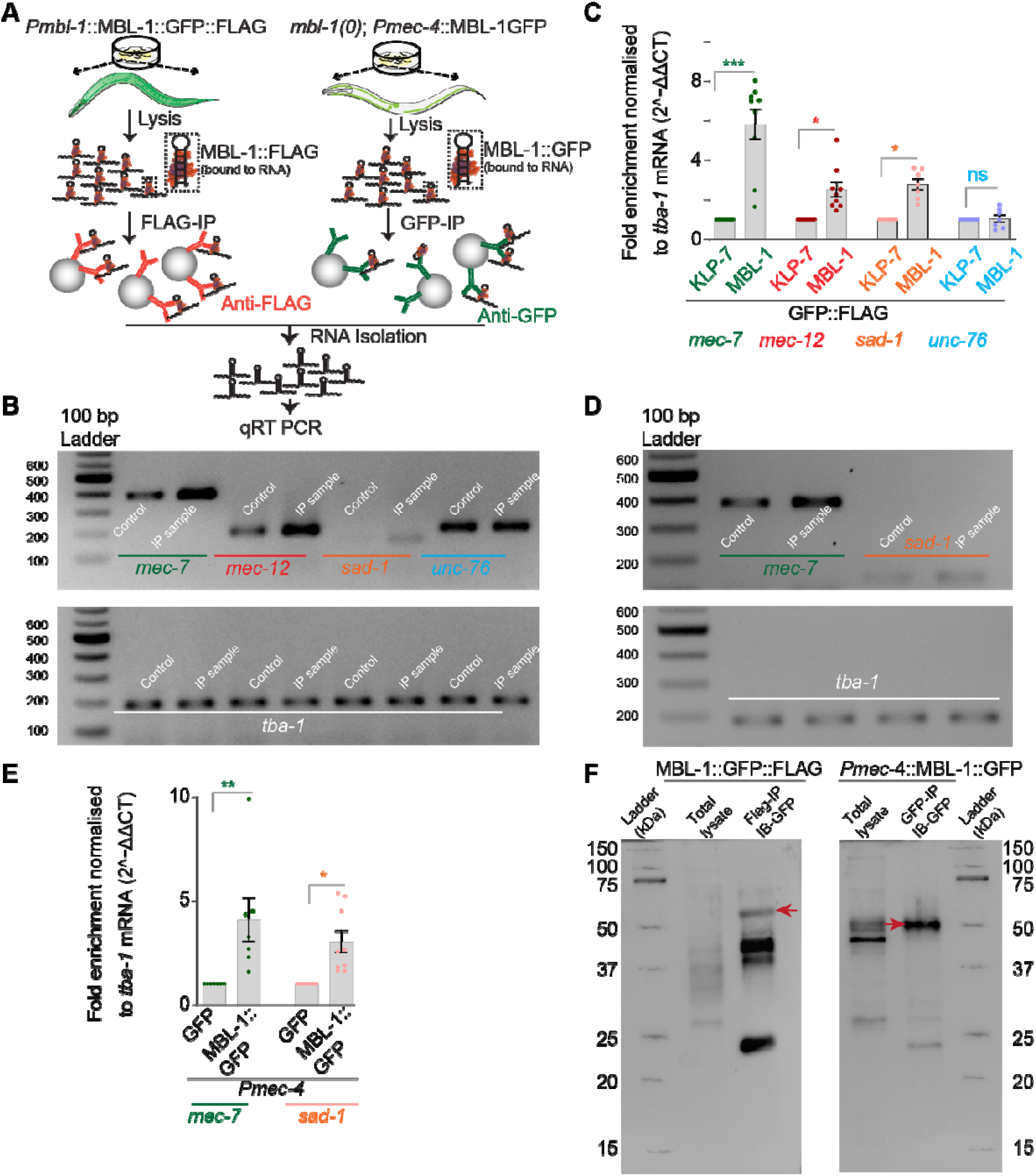
MBL-1 interacts with *mec-7* and *sad-1* mRNAs in the mechanosensory neuron. (A) Schematic illustration of Ribonucleoprotein Immuno-Precipitation (RIP-Chip) method and quantitative real-time (qRT) PCR from *Pmbl-1*::MBL-1::GFP::FLAG and *mbl-1(0); Pmec-4*::MBL-1::GFP background. (B-C) Representative agarose gel picture (B) and quantification of the transcript (C) of *mec-7, mec-12, sad-1*, and *unc-76* from the control sample (*Pklp-7*::KLP-7::GFP::FLAG) and immunoprecipitation (IP) sample (*Pmbl-1*::MBL-1GFP::FLAG). *tba-1* was used as an internal control. independent replicates (N) = 7-9 and the number of reaction (n) = 7-9. (D) Showing the representative agarose gel picture of the transcript of *mec-7* and *sad-1* from qRT-PCR from the control *mbl-1(0); Pmec-4*::GFP (*shrEx481*) and IP sample *mbl-1(0) Pmec-4*::MBL-1GFP (*shrEX75*). (E) Bar graph showing the quantification of the transcript of *mec-7* and *sad-1* from control (*mbl-1(0); Pmec-4*::GFP *(shrEx75))* and IP sample *(mbl-1(0); Pmec-4*::MBL-1::GFP (*shrEX*481). (F) Representative western blot picture showing the enrichment of MBL-1::GFP::FLAG and MBL-1::GFP (marked in red arrowhead) in the IP sample as compared to the control sample.

### Expression of *mec-7* and *sad-1* rescues the ‘short axon’ and ‘ectopic synapse’ phenotype respectively in PLM neuron

As we observed that the *mbl-1(0)* mutant has a short anterior neurite phenotype and reduced amount of total *mec-7* transcript in mechanosensory neurons due to diminished stability. We speculated that enriching *mec-7* transcript in touch neurons in the *mbl-1(0)* mutants could suppress its short anterior neurite phenotype. Indeed, when *mec-7* cDNA was expressed under its native promoter in *mbl-1(0)* mutant, the fraction of PLM neurons with short anterior neurite dropped to 20% from 80% in the *mbl-1* mutant alone (Fig. 7A-B). Consistent with this observation, overexpression of *mec-7* in *mbl-1(0)* mutants using a genomic *mec-7* construct, also significantly suppressed this phenotype. Moreover, overexpression *of mec-*7 in the wild type using the same transgene resulted in overgrowth of the anterior neurite of PLM (Fig. 7A-C).

**Figure 7:**
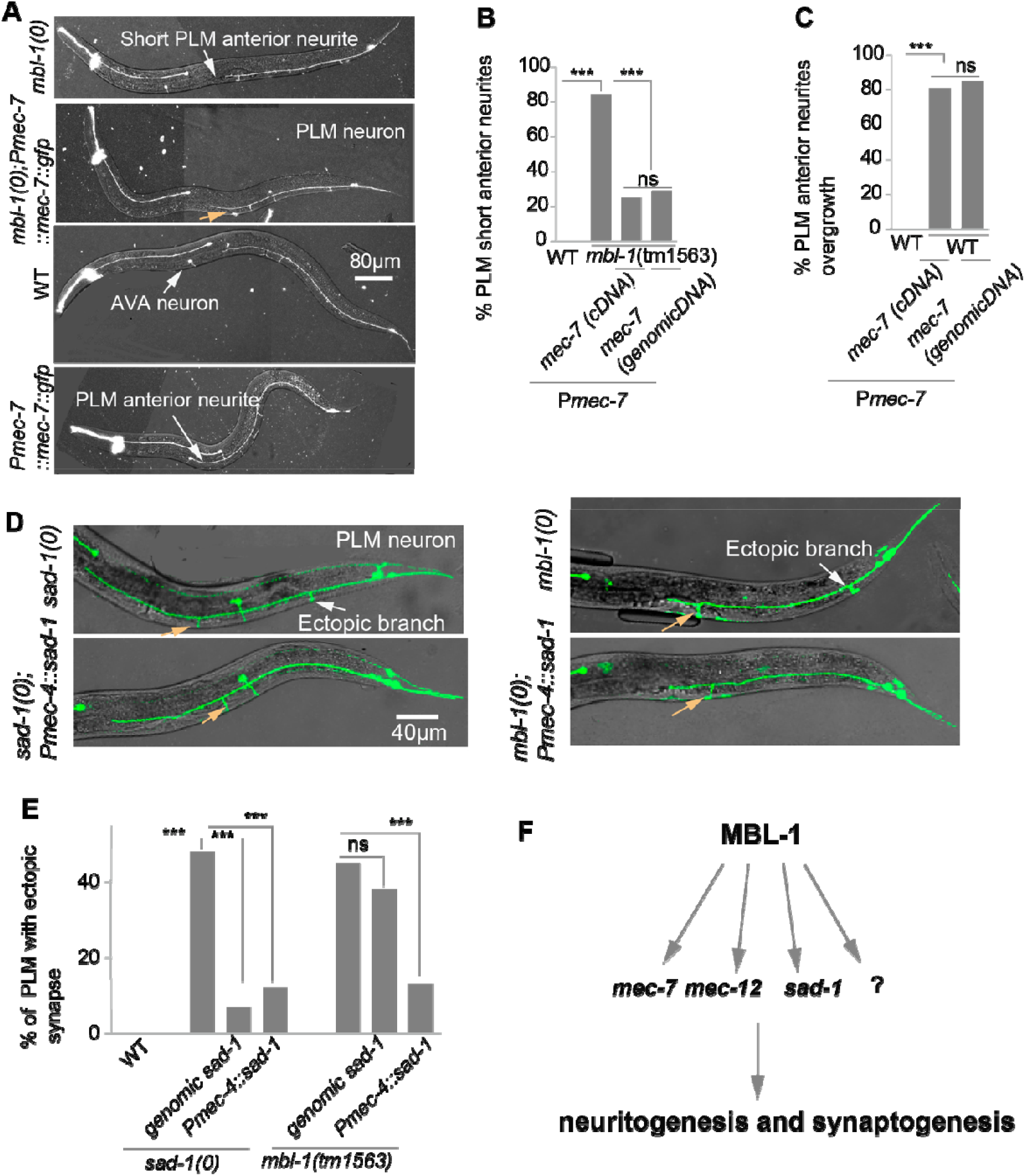
Expression of *mec-7* and *sad-1* in the *mbl-1(0)* background rescues short neurite and ectopic synapse phenotype respectively. (A) Representative confocal images of *mbl-1(0), mbl-1(0); Pmec-7::mec-7::gfp (shrEx475)*, wild-type and *WT Pmec-7::mec-7::gfp (shrEx475)*. White arrow marked in *mbl-1(0)* and WT *Pmec-7::mec-7::gfp (shrEx475)* are showing short PLM anterior and overgrowth of PLM anterior respectively. Whereas the peach arrow shows the rescue of short PLM anterior neurite. (B-C) (B) Histogram is showing quantification of short PLM anterior neurite in the different genetic backgrounds whereas (C) Histogram is showing the overgrowth of PLM anterior process in the *Pmec-7::mec-7gfp* cDNA *(shrEx475)* and *Pmec-7::mec-7gfp genomic* DNA *(shrEx476)* in the wild-type background. independent replicates (N) = 3-5 and the number of neurons (n) = 100-145. (D-E) (D) Representative confocal images showing ectopic synapse and rescue of the ectopic synapse. The bar show (E) shows quantification of ectopic synapse in *sad-1(0), sad-1(0)*; *Pmec-4::sad-1 (shrEx478), mbl-1(0)* and *mbl-1(0); Pmec-4::sad-1 (shrEx480)* background. (F) Illustration showing the regulation of *mec-7, mec-12*, and *sad-1* mRNAs by MBL-1 RNA binding protein.

We did not observe any detectable change in the stability or the amount of *sad-1* transcript in *mbl-1(0)* mutants. But we did observe an enrichment of *sad-1* transcript in MBL-1 pull-down IP samples. It was already known that MBL-1 prevents the exclusion of one exon of *sad-1* transcript in ALM touch neurons so that only the exon-included isoform is expressed (60). Hence, we speculated that the ectopic synapse defect in *mbl-1(0)* is because of a lack of exon-included isoform of *sad-1* in the *mbl-1* mutant (Figure 7D). This defect was significantly rescued by expressing the exon-included isoform in the *mbl-1* mutant using a touch neuron-specific promoter (Figure 7D-E). However, the transgenic expression of the genomic *sad-1*, in the *mbl-1* mutant fails to rescue the ectopic synapse defect (Figure 7D-E). This further strengthens the point that MBL-1 might be involved in the isoform-specific regulation of *sad-1* transcript.

## Discussion

In this work, we identified *mbl-1* mutant as a suppressor of *klp-7* mutant in which microtubules are hyper-stabilized. We found that *mbl-1* acts cell-autonomously to regulate both neurite growth and synapse formation in PLM touch neuron. Further investigation revealed that microtubule stability in PLM touch neuron is compromised in *mbl-1* mutant due to reduced *mec-7* and *mec-12* tubulin levels. We showed that MBL-1 binds to the *mec-7* and *mec-12* transcripts and regulates their stability (Figure 7F). Additionally, MBL-1 regulates the correct location of synapse in TRN by regulating the *sad-1* kinase (Figure 7F).

### MBL-1 regulates axon growth and synapse formation in neurons

RNA binding proteins (RBP) play important roles in different developmental stages of neurons, including neurogenesis, migration, pathfinding, synapse formation, axon, and dendrite growth (63, 64). However, the role of Muscleblind-1, in nervous system development is less explored. MBL-1 regulates alternative splicing, localization, stability, and processing of mRNAs (21-23). Our study reveals that the *C. elegans* homolog of MBNL-1, MBL-1, regulates the neurite growth of PLM neurons. We also observed defects in neurite growth in the BDU interneuron. In *Drosophila*, MBL-1 controls axon guidance by regulating alternative splicing of Dscam-2, cell-autonomously (37). Our data is in consistence with the role of cytoplasmic form of MBNL-1 in neurite extension in the primary culture of mouse hippocampal neurons (36). A previous study in worms showed that MBL-1 regulates synapse formation in DA9 motor neuron (45). We also observed that MBL-1 regulates synapse formation in PLM neuron. Moreover, we observed large gaps in the dorsal cord region in the *mbl-1* mutant, using the reporter for GABAergic motor neurons, which is an indication of loss of neuromuscular synapse. In the worm, similar to MBL-1 there are four other, known RNA binding proteins, MEC-8/RBMPS, MSI-1/MSI-2, UNC-75/CELF5, and EXC-7/ELAVL4, which control the splicing, stability, and localization of the transcript (65). But the loss of function mutants of these genes do not have any morphological defects in the mechanosensory neurons as seen in the *mbl-1* mutant. MEC-8 specifically regulates the alternative splicing of *mec-2* in mechanosensory neurons and controls gentle touch sensation (66, 67). However, we did not observe any morphological defect in PLM neuron due to loss of *mec-8*. It is known that Duchenne muscular dystrophy (DMD) patients show symptoms of progressive muscle degeneration in addition to learning disabilities, impaired cognitive function, and memory impairment (68, 69). DM1 patients show an age-related decline in frontotemporal functions, including memory impairment (68, 70). We noticed that GFP intensity becomes very dim in older worms. Even in L4 stage in *mbl-1* mutant, GFP intensity is dimmer (Figure 2A). This might indicate an accelerated neuronal aging of touch neurons in *mbl-1(0)*.

### MBL-1 regulates neuronal microtubule cytoskeleton for controlling axon growth and synapse formation

We isolated the mutation in *mbl-1* in a genetic screen designed to identify the regulators of microtubule cytoskeleton. Although MBNL is an RNA-binding protein, its involvement in microtubule cytoskeleton regulation and RNA transport is emerging (38, 71). A careful investigation of neuronal microtubule dynamics in *mbl-1* mutant revealed that both microtubule stability and orientation are compromised in the anterior neurite of PLM neuron. This was associated with the severe reduction in the anterograde transport of synaptic protein RAB-3. This is consistent with the previous finding that knockout of MBL-1/2 in mouse cortical neurons leads to a reduction in the dendritic complexity, and alterations in postsynaptic densities (72). These axonal and dendritic morphological changes have been linked to cytoskeletal machinery (73, 74). By combining in-silico approaches along with immunoprecipitation and RT-PCR analysis, we found that MBL-1 binds to both the transcripts of α (*mec-12*) and β-tubulin (*mec-7*) and regulates their transcript levels in touch neuron. Expression of tubulin in the *mbl-1* mutant is sufficient to restore the neurite-growth defect. However, we did not observe splicing defects in the transcripts of these genes in the absence ofMBL-1 protein. This conclusion was validated by two observations: firstly, we did not find any change in the length of the *mec-7* and *mec-12* transcripts in *mbl-1(0)* mutants, and secondly, overexpression of either *mec-7* cDNA or genomic DNA could suppress the short neurite phenotype in *mbl-1(0)*. Our data suggests that MBL-1 is regulating the stability of α (*mec-12)* and β (*mec-7)* tubulin transcripts for axon growth in the PLM neuron. However, it’s not clear how the stability of the *mec-7* or *mec-12* transcript is regulated by MBL-1. MBL-1 is known to act as an adaptor for mRNAs for their correct localization (23, 75). It’s possible that the stability of these transcripts is achieved through such mechanisms.

It is already shown that MBL-1 regulates the splicing of *sad-1* in the ALM and BDU neurons of *C. elegans* (60). The function of SAD-1 is well established in synapse formation and stabilization (6, 76). We observed ectopic synapse formation in both *mbl-1(0)* and *sad-1(0)* mutants, which indicates that these two genes could be acting in same pathway for proper synapse formation. There are two isoforms of *sad-1* and each isoform has a cell-specific expression pattern (60). When we over-expressed the exon-three included isoform in the touch neuron in the *mbl-1(0)* background, it suppressed the ectopic synapse phenotype in this background. However, over-expression of genomic *sad-1* failed to suppress the ectopic synapse phenotype. This result supports the hypothesis that MBL-1 regulates the splicing of *sad-1* to control synapse formation.

In summary, our finding shows that the RNA binding protein, MBL-1, regulates the cytoskeleton-related genes for proper axon growth and synapse formation. MBL-1 interacts with touch neuron-specific tubulin mRNAs and stabilises them for optimizing microtubule stability in PLM, which in turn promotes vesicle transport. For making the correct number of synapses at the correct location, *mbl-1* interacts with *sad-1* kinase. This study provides mechanistic insight into how an RNA-binding protein regulates the structure and function of the neuron through cytoskeletal machinery.

## Materials and methods

### *C. elegans* genetics

*C. elegans* strains were cultured on standard Nematode Growth Medium (NGM) plates seeded with OP50 *Escherichia coli* bacteria at 20°C (77). All the loss of function mutant alleles are denoted as “*0*”. For example, the *mbl-1(tm1563)* mutant is presented as *mbl-1(0)*. The wild-type N2 Bristol strain was used for removing background mutation and CB4856 Hawaiian isolates for restriction fragment length polymorphism (RFLP) mapping. The list of all the mutant and transgenic reporter strains used in this study is mentioned in Table S5 and Table S6 respectively. The extrachromosomal DNA carrying newly generated transgenes used in this study is mentioned in Table S6. Transgenes were introduced into the various mutant backgrounds by crossing. PCR or sequencing methods were used for confirming the homozygosity of all the mutants. We have used the following published transgenes in this work, *Pmec-7*-GFP (*muIs32*), *Pmec-4*-EBP-2::GFP (*juIs338)* (78), *Pmec-4*-mCherry (*tbIs222)*(79), *Pmec-7*-GFP::RAB3 (*jsIs821)* (47), and *Pmec-7*-TagRFP::ELKS-1 (*jsIs1075*) (48).

### Mapping of *ju1128* mutation by restriction fragment length polymorphism (RFLP)

For mapping of *ju1128* mutation, we crossed the *klp-7* suppressor strain bearing *ju1128* mutation (*klp-7(0); ju1128*) with a polymorphic *C. elegans* strain, the Hawaiian strain (CB4856). Individual worms from the F2 progeny of this cross were selected and single-selfed. The progeny of these F2s was genotyped for *klp-7(tm2143)* deletion mutation using forward and reverse primers. The F2 plates with confirmed genotype of *klp-7(tm2143)* deletion mutation were phenotyped for ALM ectopic extension suppression. 30-50 individual F2 plates which were homozygous for *klp-7* deletion mutation and suppression of ALM ectopic extension, were further selected for mapping. For mapping chromosomal location, DNA from these 30-50 unique F2 recombinants was pooled, and using three primers per chromosome and two primers for X-chromosome, SNP mapping was done as described in earlier protocols (43). SNP mapping result is presented in Figure S1A. As the EMS suppressor screening was done in the N2 (Bristol) strain, we used N2 SNPs to establish the linkage. In the SNP mapping gel picture, linkage of N2 SNPs with two chromosomes, the 3^rd^ chromosome, and the X chromosome was observed (marked with red arrowhead S1A). We knew that the *klp-7* gene is present on the 3^rd^ chromosome, so the 3^rd^ chromosome linkage was inferenced to be due to *klp-7* mutation. The X chromosome linkage was likely due to *ju1128* mutation (S1 marked with red arrowhead), which was consistent with whole-genome sequencing data analysis results (see below).

### Whole-genome sequencing analysis for *ju1128* mapping

*klp-7* suppressor strain, *klp-7(0); ju1128* were crossed to the Hawaiian strain (CB4856) for generating F2 recombinants. We used the same F2 recombinants which we used for SNP mapping. We prepared genomic DNA by phenol-chloroform extraction and ethanol precipitation method. We sent samples for sequencing on an Illumina HiSeq4000 platform using 50 bp paired-end reads at the core facility of the University of Washington (USA). After filtering out low-quality reads, 300 million reads were recovered, resulting in an 18X average coverage of the genome from this data. We aligned reads to the *C. elegans* reference genome version WS220 and analyzed them using the CloudMap pipeline (44). From the two aligned files, we obtained a single file of all the variations using genome-wide variant call statistics. The background variations of the parental strain (*klp-7(0*)) as well as other sister mutants isolated in the same screen, such as ju1130, were subtracted from the list of total variants, and a filtered list of candidate mutations was obtained. This list was then annotated using the available reference annotation file of *C. elegans*. For the *ju1128* mutation, we got 8 candidate genes with the *mbl-1* gene as one of them. We tested all of the candidate genes by injecting a fosmid expressing the wild-type copy of the gene into *klp-7(0); ju1128* background (Figure 1A). We observed that the fosmid-bearing an *mbl-1* gene copy, rescued suppression of *klp-7(0)* multipolar phenotype (Figure 1A).

### Widefield fluorescence imaging of mechanosensory neurons for quantifying developmental defects

Phenotyping of touch receptor neurons (TRNs) was done at the L4 stage of worms, using a Leica DM5000B microscope with a 40X objective (NA 0.75). For immobilizing worms 10 mM levamisole (Sigma-Aldrich; L3008) on 5% agarose was used. The morphology of ALM and PLM neurons was qualitatively scored using this method. This method allowed us to judge the anatomical defects in ALM neurons due to loss of *klp-7*(Figure 1A; C) or in PLM neurons due to loss of *mbl-1* (Figure 2A-B). This method was also used to quantify ectopic synapse or more than one synapse phenotype in the *mbl-1(0)* background (Figure 2C-E).

### Image acquisition and analysis of neurite length of mechanosensory neurons using a point-scanning confocal microscope

Imaging of ALM/PLM was done at the L4 stage of worm using a Zeiss Axio Observer LSM 510 Meta confocal microscope. GFP reporter, *muIs32* (*Pmec-7*::GFP) was imaged at 66 % of a 488-nm laser under a 40X oil objective (NA 1.3). ALM/PLM neurite length was normalized with respect to body length. For this purpose, differential interference contrast images were taken simultaneously with the fluorescence images.

The absolute length of the anterior and the posterior processes of PLM and the anterior process of ALM was calculated using Zeiss LSM Image Browser software or ImageJ. For PLM, anterior and posterior processes, segmented traces were drawn for getting the value of the length of these processes. The length of the anterior process of PLM was normalized with the distance between the PLM cell body and the vulva of the respective worm measured from the differential interference contrast images (Figure 4D). And the length of the posterior process of PLM was normalized to the distance between the PLM cell body and the tip of the tail (Figure 4 E). Similarly, the length of the anterior process of ALM was normalized to the distance between the vulva to the tip of the head (Figure 4F). As described earlier for touch receptor neuron length quantification (80).

### Image acquisition for GFP::RAB-3, *Punc-86*::GFP, *Punc-25*::GFP, ELKS-1::TagRFP, and UNC-9::GFP using a point-scanning confocal microscope

For imaging of GFP::RAB-3, *Punc-86*::GFP, *Punc-25*::GFP, ELKS-1::TagRFP, and UNC-9::GFP, L4 stage worms were imaged using a Nikon A1HD25 confocal microscope under 60X oil objective (NA 1.4). For GFP reporter imaging, 7 % of the 488-nm laser was used while TagRFP and mCherry were imaged using 0.3 % of 561-nm lasers.

### Molecular cloning and generation of new transgene

For making pan-neuronal, touch neuron-specific, and muscle-specific expression gateway entry clones of mbl-1 transgene (Thermo Fisher Scientific; K2500-20), the MBL-1 cDNA was single-site LR recombined with pCZGY66 (*Prgf-1* destination vector), pCZGY553 (*Pmec-4* destination vector), and pCZGY61 (*Pmyo-3* destination vector) respectively, using LR clonase enzyme (Invitrogen;11791-020). pNBRGWY29 (MBL-1 PCR8) entry clone was used for the expression of the MBL-1c isoform. To make *Pmec-4::sad-1* (pNBRGWY164), the entry clone pNBR58 corresponding to *sad-1a* cDNA was recombined with pCZGY553 (*Pmec-4* destination vector). *sad-1* was cloned into PCR8 vector using the following primers 5’ TCCGAATTCGCCCTTCGTCAATCGGGCAAAGTC 3’ and 3 ’GTCGAATTCGCCCTTGATGATAGATTAGACTTTATCAGCC 5’ with help of infusion reaction (Takara, 638947). For making *Pmec-7::mec-7::gfp* (cDNA) (pNBR165) and *Pmec-7::mec-7::gfp* (genomic) (pNBR166), we made destination vector *Pmec-7::*GWY::GFP (pNBR61) using following primers to amplifying *mec-7* promoter, 5’ CCATGATTACGCCAATGGCGCGCCAAATGTAAACC 3’ and 3’ TGGCCAATCCCGGGGCGAATCGATAGGATCCACGATCTCG 5’, and for amplifying vector backbone we used following primers, 5’CCCCGGGATTGGCCAAAG 3’ and 3’ TTGGCGTAATCATGGTCATAGCTG 5’. An infusion reaction was used to make this destination vector. We cloned *mec-7* cDNA and *mec-7* genomic DNA into the PCR8 backbone using infusion reactions. Following primers were used for cloning *mec-7* cDNA and genomic DNA: 5’ TCCGAATTCGCCCTTATGCGCGAGATCGTTCATATTC 3’ and 3’ GTCGAATTCGCCCTTCTCTCCGTCGAACGCTTC 5’. Next, these entry clones *mec-7* cDNA (pNBR59) and *mec-7* genomic (pNBR60) were recombined with pNBR61 (*Pmec-7::*GWY::GFP destination vector) to generate *Pmec-7::mec-7::gfp* (cDNA) (pNBR165) and *Pmec-7::mec-7::gfp* (genomic) (pNBR166). These plasmids were injected at different concentrations as described in Table S6. The concentrations of coinjection marker, *Pttx-3*::RFP, used was 50 ng/μl. The injection mixture’s total DNA concentration was kept at around 110–120 ng/μl by adding pBluescript (*pBSK*) plasmid to the injection mixture.

### Live imaging of EBP-2::GFP, and GFP::RAB-3 using spinning disk confocal microscopy

A Zeiss Observer Z1 microscope equipped with a Yokogawa CSU-XA1 spinning disk confocal head and a Photometric Evolve electron-multiplying charge-coupled device camera for fast time-lapse image acquisition was used for EBP-2::GFP and GFP::RAB-3 imaging. For EBP-2::GFP, images were taken at 2.64 frames per second for a total of 2-minutes duration. For GFP::RAB-3, images were taken at 3.19 frames per second for 3 minutes. To get the best signal-to-noise ratio for EBP-2::GFP, 8.75mW of a 488-nm excitation laser was used while for GFP::RAB-3 10mW power was used.

### Analysis of EBP-2::GFP and GFP::RAB-3 dynamics

The kymographs of EBP-2::GFP (Figure 3B) and GFP::RAB-3 (Figure 3H) were generated using the Analyze/ MultiKymograph tool in ImageJ software (https://imagej.nih.gov/ij/) from 30-μm ROIs placed on the PLM anterior and posterior process (Figure 3A). In both types of kymographs, the horizontal axis is representing the axon length in micro-meters, and the vertical axis represents time in seconds. For EBP-2::GFP movies, ROIs were drawn in distal to proximal direction (towards the cell body) for the anterior process of PLM and proximal to distal for the posterior process of PLM. The diagonal tracks which are moving away from the cell body were annotated “plus-end-out” microtubules (P; green traces in Figure 3B), and the diagonal tracks which are moving toward the cell body were denoted as “minus-end-out” microtubules (M; magenta traces in Figure 3 B). The fraction polarity of microtubules was calculated from the relative number of plus-end out tracks or minus-end out tracks to the total number of tracks in a given EBP-2::GFP kymograph. The growth length and growth duration of EBP-2::GFP tracks were calculated as a net pixel shift in the X and Y axes, respectively, (Figure 3E-F).

For GFP::RAB-3 movies ROIs were also drawn similar to EBP-2::GFP movies. The diagonal tracks which are moving away from the cell body were annotated “anterograde” (green traces in Figure 3H), and the diagonal tracks which are moving toward the cell body were denoted as “retrograde” (magenta traces in Figure 3H).

We calculated anterograde and retrograde particle movement from each kymograph by quantifying the number of tracks in either anterograde or retrograde direction from 30-μm ROIs corresponding to the anterior and posterior process of PLM near the cell body during the 3 minutes of imaging. We calculated run length and run duration by quantifying a net pixel shift in the x and y axes respectively in the anterograde and retrograde directions (47).

### Gentle touch assay

The L4 stage hermaphrodite worms were subjected to a gentle touch assay using a tip of the eyelash. The worms were touched at the anterior and posterior ends alternatively, 10 times each, as discussed previously (39, 79). A response was considered positive, if it elicited a reversal behavior. We denoted a positive response as 1 and no response as 0. The anterior touch response index (ATRI) (Figure 3M) and posterior touch response index (PTRI) (Figure 3N) was calculated as a ratio of the total number of responses to the total number of touches given (10 touches).

### Identification and analysis of MBL-1 targets

To identify MBL-1 targets in the PLM neuron, we ascertained genes expressed in PLM neurons using the CenGEN database (52) for a threshold value of 2. We found that 5,283 genes are expressed in the PLM neuron (Table S1) (CenGEN database). When we put 5,283 genes expressed in PLM neuron in the oRNAment database <u>(h</u>t<u>tp://rnabiology.ircm.qc.ca/oRNAment)</u> and looked for MBL-1 binding site CGCU in these genes, we got 2000 genes with potential MBL-1 binding site and an enriched expression in PLM neuron. Using gene ontology (GO) analysis, we shortlisted genes involved in (1) Microtubules-based process (2) axon development (3) regulation of synapse structure (4) Axodendritic transport. The analysis is presented in Figure 4A, Table S2.

### Reverse transcription (RT) PCR for checking the splicing of the transcript

For checking the splicing event, we used the reverse transcription method. We collected the total RNA from wild-type and *mbl-1(tm1563)* mutant at day one adult staged worm (A1). We synchronized the worms by allowing the gravid adult worm to lay eggs for 30 minutes and then progeny was grown at 20 °C till the A-1 adult stage. The staged worms were washed thrice with M9 buffer and the worm pellet was collected and stored at -80° for RNA isolation. RNA isolation was done using Qiagen RNeasy Mini Kit (no. 74104; Qiagen) from thawed worm pellet. The extracted RNA was treated with DNase I (Ambion’s DNA-free kit AM1906) to get rid of any genomic DNA contamination. cDNA was reverse transcribed from ∼3-4 µg of this treated RNA using Superscript III Reverse Transcriptase (18080093). For checking the splicing, we used 200 ng cDNA from a wild-type and *mbl-1(tm1563)* background for a 25 μl PCR reaction. We used emerald Amp GT PCR (2X master mix, cat-RR310) Taq polymerase, and the primers which were used for checking the transcript of *mec-7, mec-12, sad-1*, and *unc-76* are given in Table S3, and primer binding site is showed in Figure S5A. For *mec-7* and *mec-12*, we have designed primers in such a way that we amplified the full length of transcript and these genes have only one isoform (Wormbase-WS285). We used 30 cycles for amplification. The 5 μl PCR products of *mec-7, mec-12*, and *aak-2* (we used as a control reaction) transcripts were run on a 1% agarose gel (Figure 5A) while *sad-1* and *unc-76* PCR products were run on 2% agarose gel (Figure S5B).

### Quantitative real-time (qRT-PCR) for checking the total transcript

We have used A1-worms for isolating the total RNA as discussed in the previous section from the wild-type and *mbl-1(0)* mutant background. The extracted RNA was treated with DNase I (Ambion’s DNA-free kit AM1906) to get rid of any genomic DNA contamination. We reverse-transcribed ∼2-3 µg DNase treated RNA into cDNA using Superscript III Reverse Transcriptase (Invitrogen no. 18080093). For each reaction, 50ng of this cDNA was added to 20ul of Power SYBR Green PCR Master Mix (Applied Biosystems Life Technologies, no. 3367659). The amplification was performed for 40 cycles. The primer sequences and relative positioning are given in the supplementary information Table S4 and S5A respectively. While designing the primers for this experiment some precautions were taken. First, the primers were selected such that a single Ct peak was obtained for each qRT PCR reaction to maintain the specificity of the amplicon being quantified. Second, to avoid any amplification from genomic DNA contaminants, we selected some primers with binding sites at the intron-exon boundary. When PCR reactions were run on a 3% agarose gel, no contaminating bands were observed. The relative mRNA amounts of target genes in the *mbl-1(tm1563)* and the wild-type N2 strains were calculated using the standard ΔΔCt method (81) and were normalized to *tba-1* as a control for endogenous mRNAs (82). We have used the ΔΔCT method for calculating the fold change of transcript in the wild-type and *mbl-1(tm153)* background (81).

### Inhibitor Experiment

The RNA synthesis was inhibited by feeding worms 400μM actinomycin-D (Sigma, A9415), dissolved in DMSO, as previously described (83, 84)(. OP-50 bacteria was cultured in B-broth media overnight at 37°C in a BOD incubator. For making 400μM Actinomycin D working concentration, we have diluted actinomycin D into OP50 B-broth. For control, we have used the same amount of DMSO. This culture was then used to seed a 60 mm NGM plate. We transferred gravid adults worm for 30 minutes for egg-laying, to age synchronize worms, on NGM plates containing actinomycin D drug and DMSO. Age synchronized A1 adult worms, grown on these plates, were then washed with 1XM9 three times, collected, and frozen at -80°C. These worm pellets were then thawed on ice. From these frozen pellets, we isolated RNA and did qRT PCR as described above.

### Ribonucleoprotein-Immuno Precipitation (RIP)

Immunoprecipitation experiments were done using protocols as previously described (85). Around 50-100 gravid adults were transferred to thirty 60mm NGM plates and allowed to lay eggs for half an hour at 20°C. The progeny was then grown till the day-1 adult (A1) stage at 20°C. These A1 synchronized worms were pooled, washed thrice with 1XM9, and pelleted at 1500 rpm for 2 minutes. The collected worm pellet, which was more than 300 µl in volume, was then stored at -80°C until further use. It was then thawed on ice for homogenization and all further procedures were carried out at 4°C. Worm pellets were homogenized in 400-500µl of ice-cold 2X lysis solution [Buffer A (20mM Tris (pH 8.0), 150 mM NaCl, 10mM EDTA) + 1.5mM DTT, 0.2% NP-40, 0.5 mg/ml Heparin Sulphate, 1X EDTA complete Protease inhibitor (Roche -11836153001, 1 tablet for 5ml), RNase inhibitor (Invitrogen AM2696, 50U/ml), Phosphatase inhibitor (100U/ml and RNase out (Invitrogen 1643272, 100U/ml)] The homogenized sample was passed successively through 19mm, 22mm, 26mm, and an Insulin syringe to make a smooth homogenate which was then centrifuged, at 19,000Xg for 20 minutes, to obtain a clear supernatant. 5-10% of the total lysate was kept aside as total input for RNA estimation and western blotting. The total protein concentration in each sample was determined by the Bradford assay. Equal amounts of protein across all conditions were used for the RIP experiments.

Agarose beads (Roche, 11719416001) were equilibrated in 1X lysis buffer. For pre-clearing, supernatants were each incubated with 20µl of equilibrated agarose beads for one hour, followed by centrifugation at 8,000Xg for ten minutes to collect the beads. The supernatant collected was the pre-cleared lysate that was further processed.

To immunoprecipitate Flag-tagged GFP from strains, anti-Flag M2 agarose beads (Sigma, A2220-1ML) were used. For each sample 60µl of 50% slurry (i.e. 30µl of packed beads) of anti-Flag M2 agarose beads was taken in a 1.5 ml tube and washed with 1X lysis buffer to equilibrate. Pre-cleared lysates from the previous step were added to the equilibrated beads and incubated at 4°C, overnight, with continuous mixing. For immunoprecipitation of *Pmec-4*::MBL-1::GFP from wild-type and *mbl-1* samples expressing this transgene, 3ug of anti-GFP antibody (MBL LifeSciences M048-3, raised in mouse) was added to the precleared lysates. and incubated for 8-10 hours with continuous mixing. The following day, 30ul of equilibrated Protein G – agarose beads (Sigma 15920010) were added to the lysates and incubated further for 3 hours.

Both samples (anti-Flag M2 and anti-GFP) were centrifuged at 10,000 rpm for 15 minutes at 4°C. The pellets were collected and washed with lysis buffer containing 150 mM NaCl, followed by centrifugation at 10,000Xg for 15 minutes at 4°C. The agarose beads were collected, 20% of the bead’s volume was kept for western Blot and the remaining 80% was used for RNA isolation and qRT PCR as described above.

### Western blot

Samples were boiled in Laemmli buffer and resolved on 12% SDS-PAGE. Post-transfer of proteins on nitrocellulose membranes, the membranes were blocked with 5% BSA for one hour, and then probed with anti-GFP antibody (Abcam ab290) overnight. Following day, membranes were washed in 1X Tris Buffer Saline containing 0.1% Tween 20 (TBST) and incubated with secondary antibody (anti-Rb) for 3 hours, followed by detection using standard ECL chemi-luminescence detection kit (Millipore, WBKLS0500).

### Image Acquisition and Quantification for *Pmec-7::mec-7*::*gfp*

The worms co-expressing *Pmec-7::mec-7::gfp* (cDNA, *shrEx473), Pmec-7::mec-7::gfp* (genomic DNA, *shrEx474*) and *Pmec-4::mCherry* (*tbIs222)* were immobilized using 10mM Levamisole. *mec-7::gfp* and *mCherry* (constitutive reporter) were imaged with 10% of 488nm for cDNA and 1.5 % for genomic DNA and 0.3% of 543 nm lasers, respectively, under a 60X oil objective (NA 1.4) of a Nikon confocal microscope. Average *mec-7::gfp* and *mCherry* intensities were measured from 50 μm anterior and posterior processes of PLM neuron and PLM cell body using ImageJ. We have placed similar ROIs outside the PLM neuron for background correction. We have plotted the ratio of GFP/mCherry (Figure 5 F) and the absolute intensities of GFP and mCherry (Figure S6 A-B).

### Statistical analysis

GraphPad Prism software version 9.0.2 was used for analyzing the data. The data presented in each figure, and the bar in the plots represents the mean value and the standard error of the mean (SEM). The X^2^ tests (Fisher’s exact test) were used for comparing the proportions. ANOVA with a post hoc Tukey’s multiple comparisons test was used for comparing more than two groups. We have used Bartlett’s test for testing the homogeneity of variances before proceeding with ANOVA. In each panel of the figure, the P-value, which is used as a measure of significance, has been presented to compare the respective group. The sample number (n) for given experiments has been presented in the respective figure legend for each graph. The total number of biological replicates (N) has been also mentioned in each figure legend of the graph.

## Abbreviations

MT: Microtubule
TRN: Touch receptor neuron

## Acknowledgments

We would like to thank the National Bio-Resource Project, Japan, and the Caenorhabditis Genetics Centre for strains. We thank Andrew Chisholm and Yishi Jin for their support and guidance at the initial stage of this project. The *ju1128* mutant was isolated in their labs. We thank Sibaram Behera for helping with touch assay experiments and for the help in Bioinformatics analysis. We also thank Sunanda Sharma for her help in editing this manuscript. We thank Arnab Mukhopadhyay, Sandhya Koushika, Michael Nonet, and Cori Bargmann for the help with strains. This work was supported by the National Brain Research Centre core fund from the Department of Biotechnology, The Wellcome Trust DBT India Alliance (IA/I/13/1/500874), and a grant from the Science and Engineering Research Board (SERB: CRG/2019/002194) to A. Ghosh-Roy. The Caenorhabditis Genetics Centre is supported by the National Institutes of Health Office of Research Infrastructure Programs (P40 OD010440).

The authors declare no competing financial interests.

## Author contributions

D. Puri, S. Samaddar, S. Banerjee and A. Ghosh-Roy designed experiments. D. Puri and S. Samaddar performed experiments and analyzed data. A. Ghosh-Roy and D. Puri wrote the manuscript.

## Notes

### Competing Interest Statement

The authors have declared no competing interest.

