## Supporting File for "Musleblind-1 regulates microtubule cytoskeleton in *C. elegans* mechanosensory neuron through tubulin mRNAs"

**List of supplementary figures and tables**

**Figure S1: Mapping of the *ju1128* mutation on the chromosome, related to Figure 1**

**Figure S2: *mbl-1* mutant display defect in the development of different classes of neurons, related to figure 2**

**Figure S3: *mbl-1* mutant displays defect in the microtubule organization in PLM neuron, related to figure 3**

**Figure S4: MBL-1 binding site in different transcripts, related to figure 4**

**Figure S5: Quantification of different transcripts in *mbl-1(0)* background, related to figure 5**

**Figure S6 Quantification of *Pmec-7*::MEC-7::GFP in *mbl-1(0)* background, related to figure 5**

Table S1: Genes expressed in PLM neuron

Table S2: GO analysis of MBL-1 targets involved in different biological processes

Table S3: MBL-1 targets checked for a phenotype in PLM neuron

Table S4: Primers used for doing qRT-PCR

Table S5: Strains used in this study

Table S6: Transgenes Generate

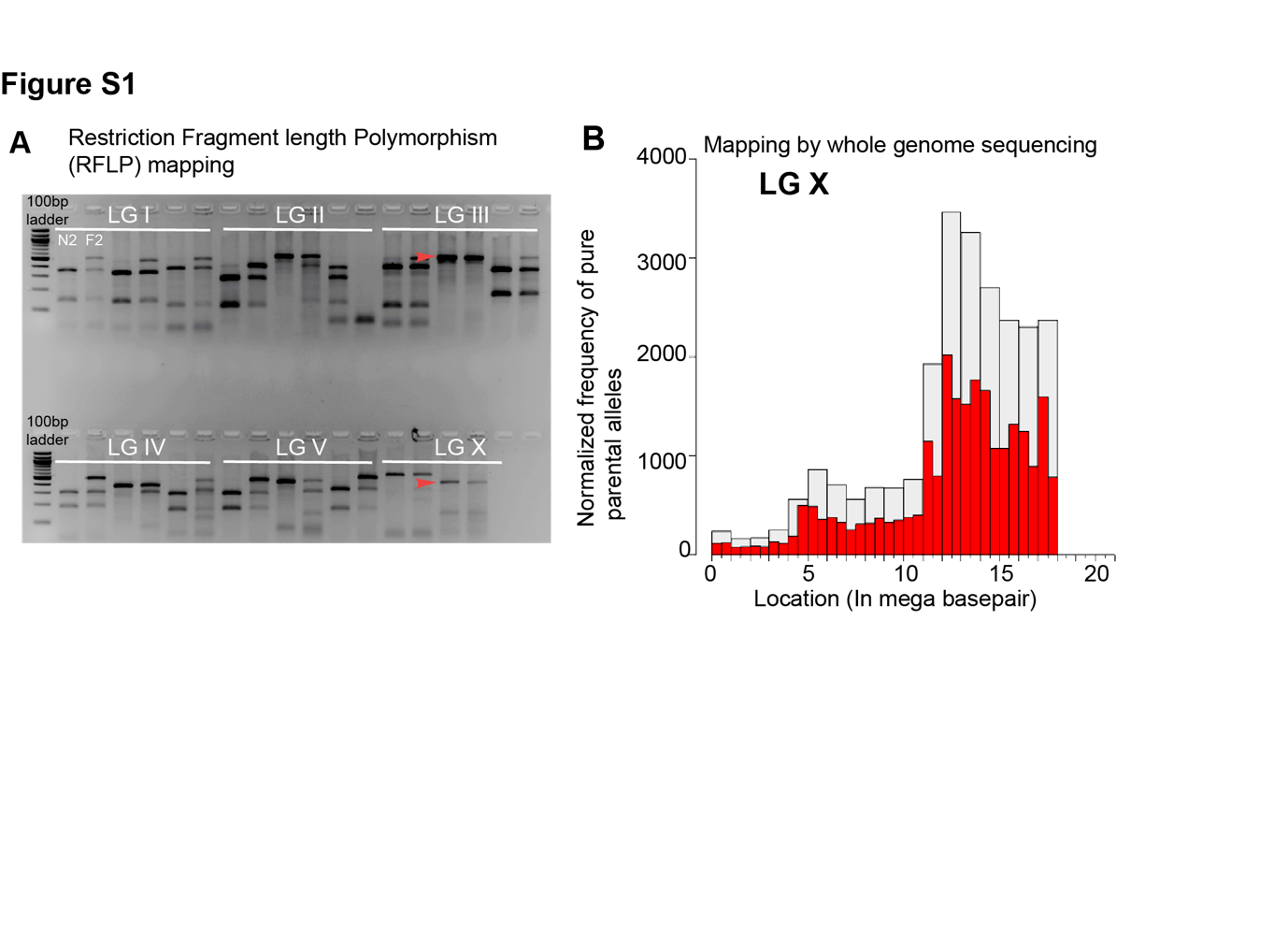

Figure S1: (A) Gel picture showing result for Restriction Fragment Length Polymorphism(RFLP) used for mapping of *ju1128* mutation. The 100-base pair ladder was used as a marker. In the gel picture, two chromosomes showed linkage- the third chromosome showed linkage for *klp-7(tm2143)* (marked with red arrowhead) and the X chromosome for *ju1128 mutation* (marked with red arrowhead). (B) The mapping of the *ju1128* mutation from the whole genome sequencing data using the method described by Minevich et al. (2012), with chromosome position (in megabases (Mb)) plotted against the Normalized frequency of pure parental alleles. *ju1128* mutation in *the mbl-1* gene is located at the 17002646 base pair position of chromosome X in which C changes to T as the result of the stop codon introduced in the *mbl-1* transcript.

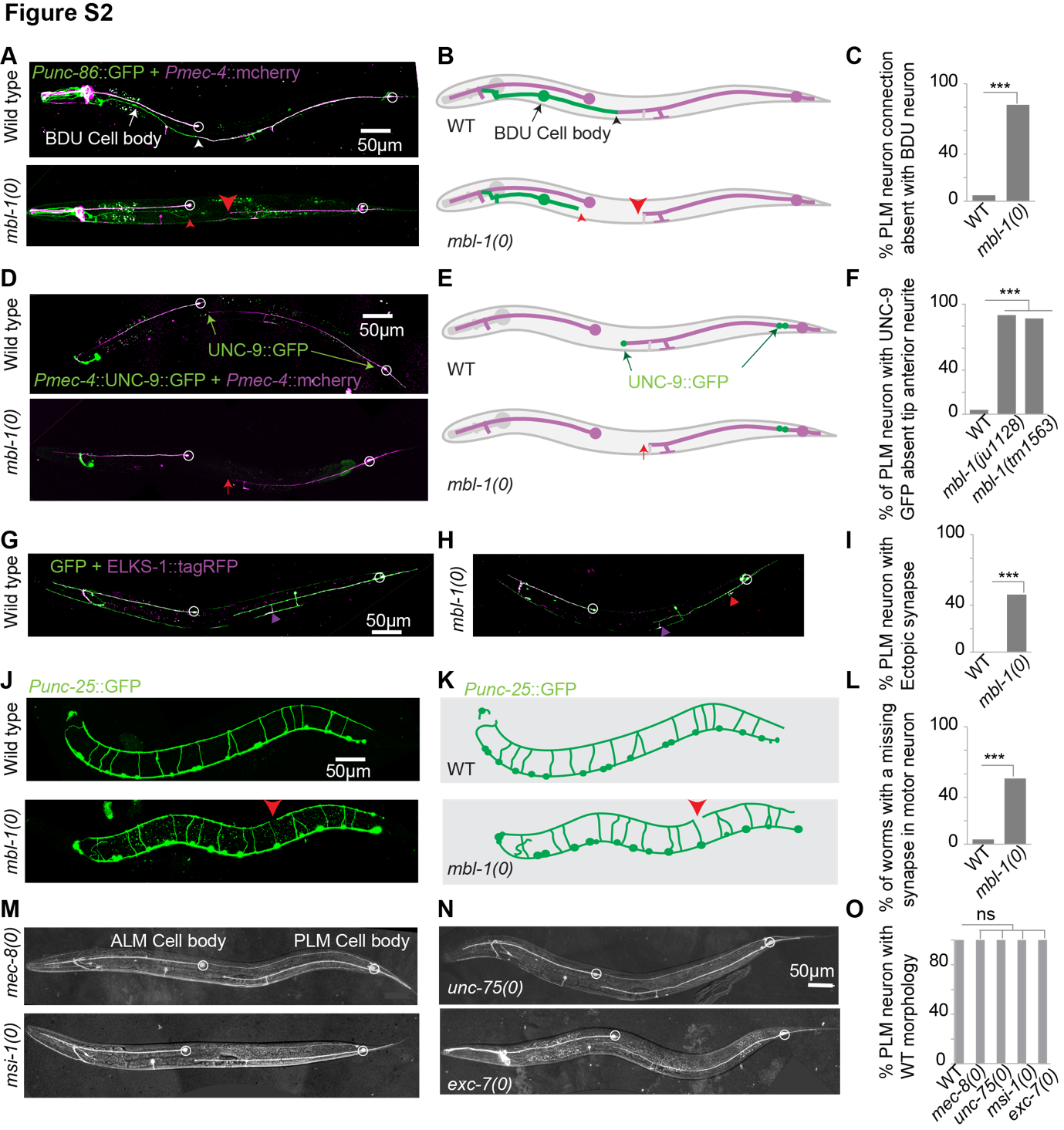

Figure S2: Images (A) and schematic (B) of touch neurons (ALM and PLM) and BDU neurons in both wild-type and *mbl-1(0)* at the L4 stage. BDU neurons were visualized with the *Punc-86*::GFP (*kyIs262*) while for visualization of touch neurons *Pmec-4*::mCherry (*tbIs222*) and *Punc-86*::GFP (*kyIs262*) transgenes were used. The presence of physical contact between PLM anterior and BDU neuron is shown by the white arrowhead in the wild-type background which is lost in *mbl-1(0)* shown by the red arrowhead. (C) Quantification of the defect as shown in the image (A). N=3 independent replicates, n (number of worms) = 25-32. (D-F) Representative confocal images (D) and schematic (E) of UNC-9::GFP in the touch neurons, in the wild-type and *mbl-1(0)* background. Green arrow showed location of UNC-9::GFP in the wild-type background whereas the red arrow points to missing UNC-9::GFP at PLM anterior tip in the *mbl-1(0)*. (F) Quantification of % of defect shown in the image (D). N=3-4 independent replicates, n (number of worms) = 30-35. (G-I). Confocal images of wild type (G) and *mbl-1(0)* (H) worms expressing *Pmec-7*-ELKS-1::TagRFP (*jsIs1075*) + P*mec-7*-GFP (*muIs32*). The ectopic synapse in the PLM anterior process in the mbl*-1(0)* background is marked in red arrowhead whereas the original synapse is marked in magenta arrowhead. (I) The histogram shows % of worms having ectopic synapses in the *mbl-1(0)* background. N=3 independent replicates, n (number of worms) = 25-30. (J-L) Representative images (J) and schematic (K) of motor neurons in the wild-type and *mbl-1(0)* at the L4 stage. Red arrow showing defect in the motor neuron in *mbl-1(0)* background. (L) The histogram shows the % of synapse defects in the motor neuron in the *mbl-1(0)*. N=3-4 independent replicates, n (number of worms) = 30-50. (M-O) Representative confocal images of RNA binding protein mutants *mec-8(0)*, *msi-1(0)* (M), *unc-75(0)*, and *exc-7(0)* (N). (O) Quantification of PLM morphology of loss of function mutants of RNA binding proteins which are shown in M-N. N=3 independent replicates, n (number of worms) = 80-130. For C, F, I, L, and O ***P <0.001; Fisher’s exact test. ns means not significant.

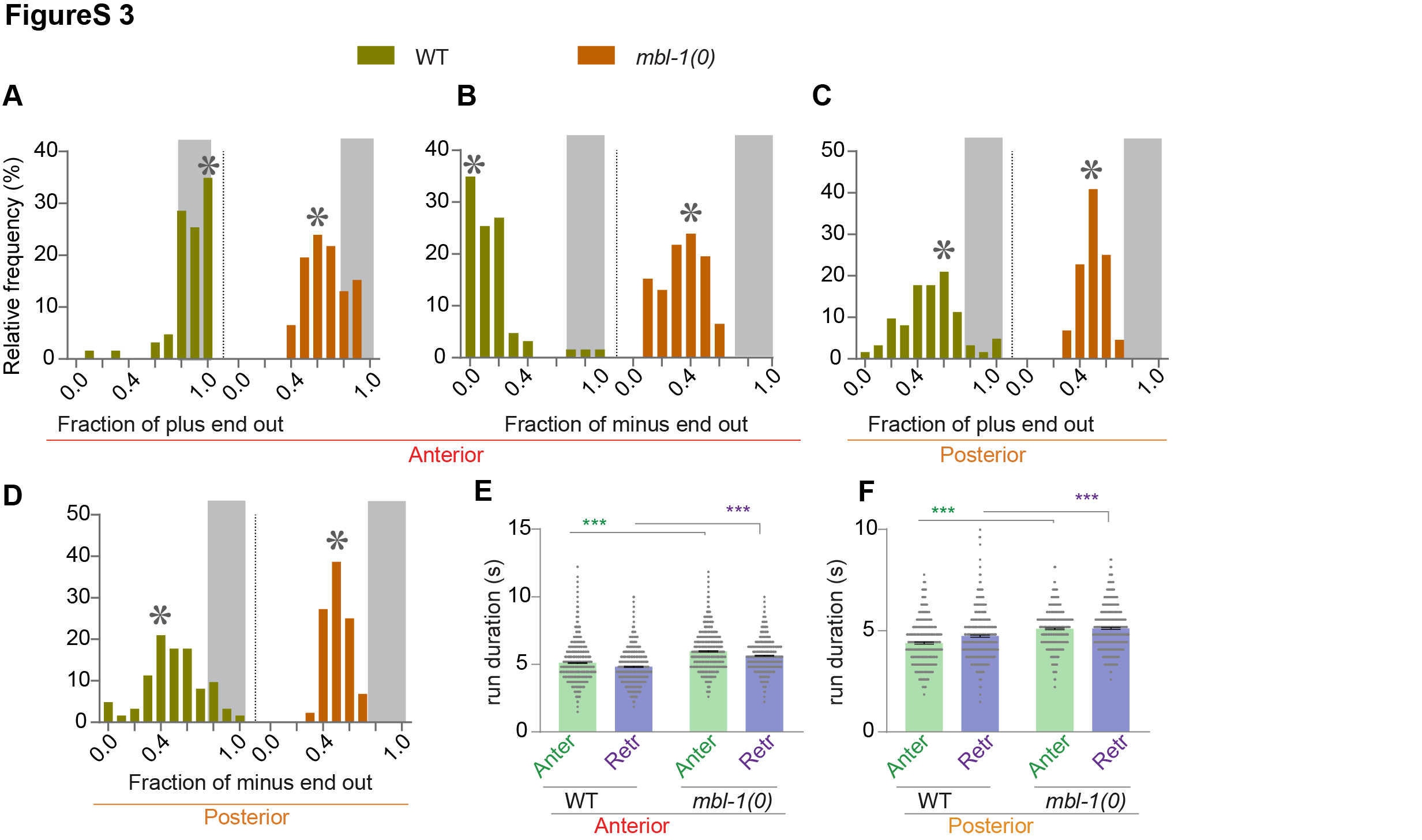

Figure S3: Frequency distribution of the fraction polarity values of microtubules with plus-end-out (A and C) or minus-end-out (B and D) microtubules organization in PLM anterior (A-B) and posterior (C-D) processes, in wild-type and *mbl-1(0)*. The mode value of each distribution is marked by the asterisk sign and the microtubules having fraction polarity value 0.8 or above are marked by a gray shaded area which represents plus-end out or minus out unipolar microtubule arrangement. N=3-5 independent replicates, n (number of worms) = 44-62. Error bars represent SEM. Statistical comparisons were done using ANOVA with Tukey’s multiple comparison test (E-F), ***p < 0.001.

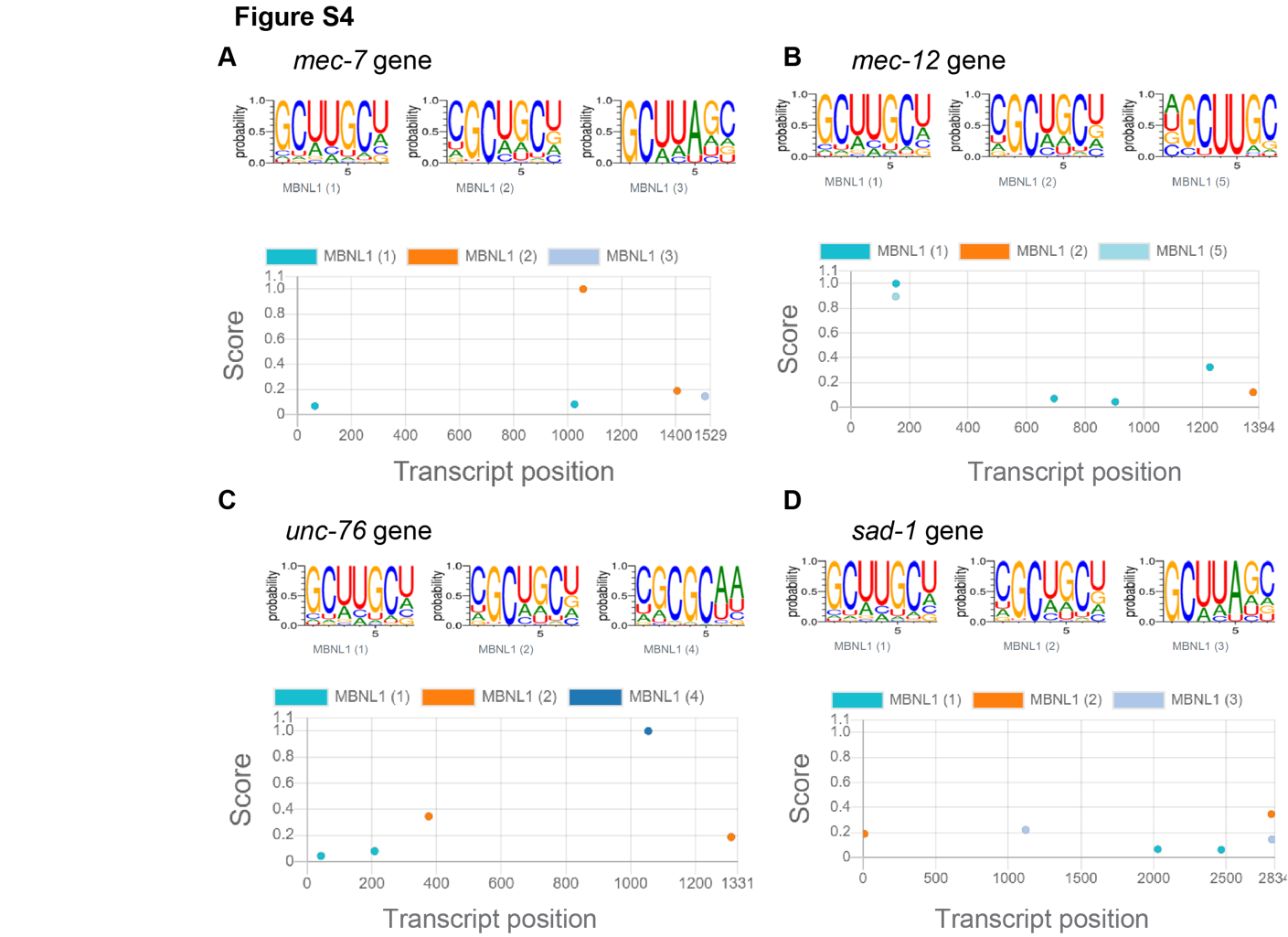

**Figure S4:** (A-B) Pictures depicting MBL-1 preferential binding sequence and binding position in the transcript of *mec-7* (A) and *mec-12* (B) genes. (C-D) Picture showing MBL-1 sequence and binding position in the transcripts of *unc-76* (C) and *sad-1* (D).

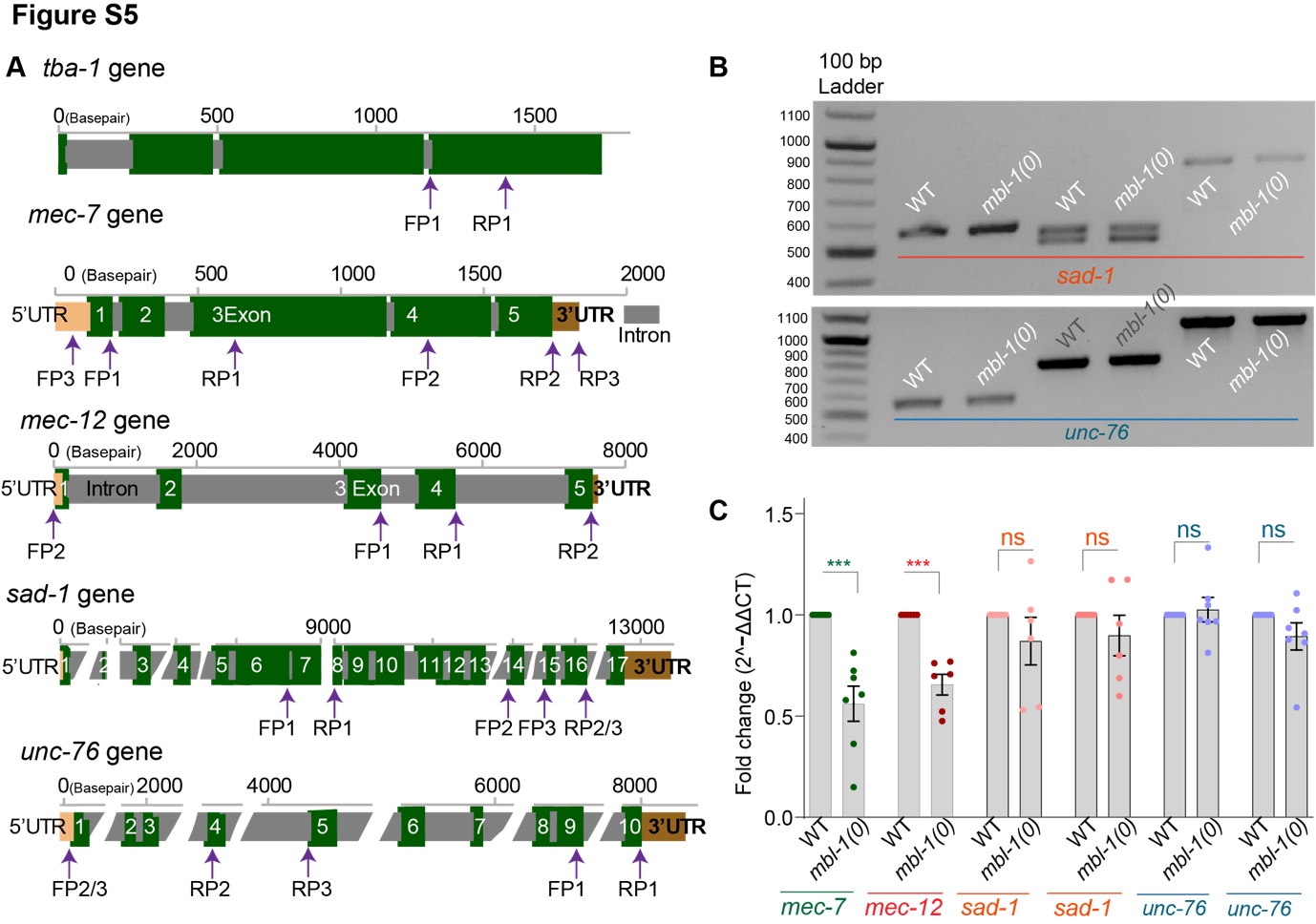

Figure S5: (A) Illustration showing the different primer positions used for checking the transcript length or doing qRT-PCR for *tba-1, mec-7, mec-12, sad-1,* and *unc-76*. The sequence of these primers is given in the supplementary file table S4. (B) Representative agarose gel image showing *sad-1* and *unc-76* transcript in the wild-type and the *mbl-1(0)* background. (C) The histogram is showing the quantification of the fold change of the transcript of *mec-7, mec-12, sad-1*, and *unc-76* in wild type and the *mbl-1(0)* background. These data were obtained from quantitative real-time PCR (qRT-PCR). independent replicates (N) = 6 and the number of reaction (n) =6-8. For ***, P < 0.001; ANOVA with Tukey’s multiple comparison test. Error bars represent SEM.

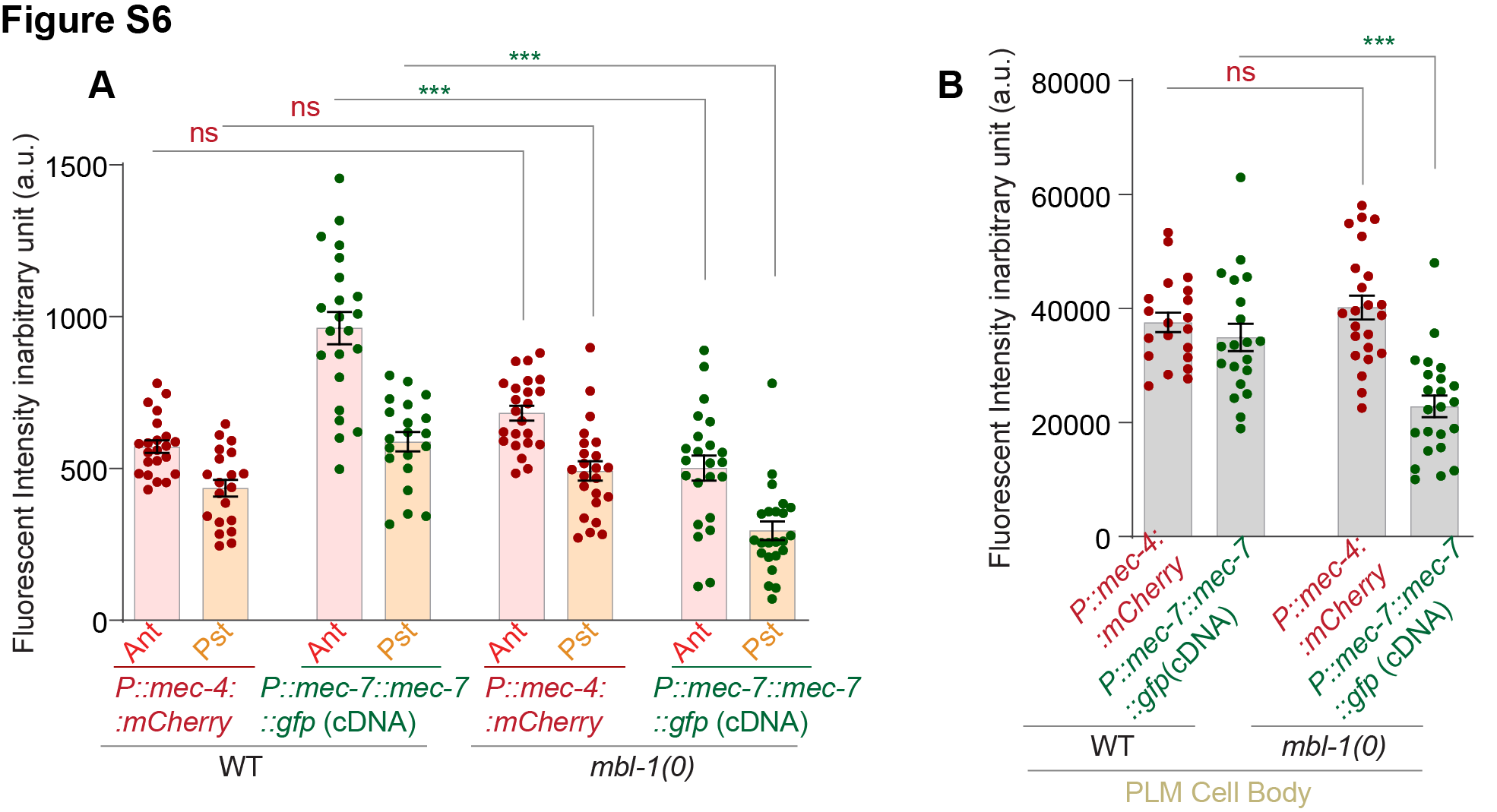

Figure S6: (A-B) Histogram showing the fluorescence intensity of *Pmec-7*::MEC-7::GFP cDNA *(shrEx473)*, *Pmec-7*::MEC-7::GFP genomic DNA *(shrEx474)*, and *Pmec-4*::mCherry in the wild-type and *mbl-1(0)* background. The fluorescence intensity is shown in the arbitrary unit in the anterior and posterior process of PLM neuron (A) from 50μm as shown in Figure 5 F and the PLM cell body (B) in the wild-type and *mbl-1(0)* background.
